# Rett Syndrome astrocytes disrupt neuronal activity and cerebral organoid development through transfer of dysfunctional mitochondria

**DOI:** 10.1101/2023.03.02.530903

**Authors:** Danielle L Tomasello, M. Inmaculada Barrasa, David Mankus, Katia I Alarcon, Abigail KR Lytton-Jean, X. Shawn Liu, Rudolf Jaenisch

## Abstract

Studies on the function of Methyl CpG binding protein 2 (MECP2) and the consequence of *MECP2* deficiency and duplication have largely focused on neurons. The function of MECP2 in human glia, along with the comprehensive understanding of glial function in neurodevelopmental disorders, is much less understood. Using female and male human embryonic stem cell (hESC) lines to model *MECP2* loss-of-function (LOF) in Rett Syndrome (RTT) in the developing brain, we investigated the molecular underpinnings of astrocyte (AST) development and dysfunction, and the mechanisms by which AST contribute to neuronal activity. Here we show that hESC-derived RTT ASTs have fewer mitochondria yet similar levels of reactive oxygen species compared to isogenic controls (CTR). We identified significantly diminished mitochondrial respiration that was compensated by increased glycolysis, and that the molecular mechanism behind mitochondrial dysfunction were reduced key proteins around the tricarboxylic acid (TCA) cycle and electron transport chain (ETC). We found an increased abundance of cytosolic amino acids in RTT ASTs under basal conditions that was readily depleted when energy demands were increased. We determined that RTT AST can donate their mitochondria to hESC-derived cortical neurons, and that isolated mitochondria from RTT ASTs are sufficient to cause significant changes to neuronal activity, increasing local field potentials to a hyperexcitable state. To examine mitochondrial health in the developing brain, we derived cerebral organoids. Ultrastructural analysis indicated that mitochondria from RTT hESC-derived organoids were significantly smaller compared to mitochondria from CTR organoids, indicating decreased connectivity and function, and this phenotype was stronger in glia compared to neurons. Using a multiomics epigenetics approach, we found hallmarks of RTT developmental delay and glial specific gene expression changes that corroborate altered energy metabolism and mitochondrial dysfunction. Based on these results, we propose that release of dysfunctional mitochondria from RTT ASTs to neurons furthers pathophysiology of the syndrome.

## Main

Rett syndrome (RTT) is a devastating neurodevelopmental disorder. Classical RTT diagnosis, most commonly due to mutations in *MECP2*, is associated with severe mental disability and autism-like syndromes that manifests during early childhood. MECP2 is a pleiotropic DNA- and RNA-binding protein and multifunctional regulator of chromatin remodeling, gene expression and other cellular pathways that remain topics of intense study^1^. This protein is expressed in all cells of the brain, yet work is still ongoing to determine the wide neurological changes due to *MECP2* LOF in RTT^2^.

ASTs provide structural and molecular support for neurons that is essential for proper development and maturation^3^. Dysfunction in ASTs has been commonly observed, largely in mouse models of RTT. Mecp2 deficiency in ASTs resulted in decreased expression of glutamate transporters EAAT1/2 and increased expression of glutamine synthetase, which together resulted in a high concentration of extracellular glutamate and excitotoxicity for neurons^4^. Increased current from GABA transporter 3 was observed in *Mecp2 KO* mouse hippocampal ASTs^5^. Adverse effects to morphology and function in neurons were observed due to AST impact and their conditioned media in both RTT mouse model and patient induced pluripotent stem cell RTT models^6–8^. Further, mouse studies have demonstrated that loss of *Mecp2* function in ASTs is sufficient to cause RTT phenotypes, and that AST-specific restoration of Mecp2 expression ameliorates these same phenotypes, highlighting the non-cell-autonomous contribution to RTT pathophysiology^9^.

Recently, a group studied a RTT causing mutation MECP2 R270X in a male hESC line^10^. They identified hESC-derived ASTs show increased glycolysis, and mitochondrial respiratory defects with basal and maximal oxygen consumption. After identifying changes in morphology when ASTs and neurons were co-cultured and in neuron spheroids, they hypothesized that dysfunctional mitochondria may impact neuronal function and maturation. What remains elusive are the molecular and biochemical changes in ASTs due to MECP2 LOF that underlie AST dysfunction and contribute to human neuron RTT pathophysiology. We aimed to define these changes by studying human stem cell-derived ASTs and their interaction with neurons in a 2D and 3D context. Corroborating findings from Sun et al.^10^, we found dysfunctional mitochondria and altered energy metabolism in RTT AST. Here, we defined the molecular mechanism of mitochondria dysfunction through integration of metabolomic and proteomic analysis on isolated AST mitochondria. We provide new novel insight into RTT pathology after we determined the cellular mechanism by which dysfunctional mitochondria from RTT ASTs impact neuron function, and found glial specific mitochondrial and metabolic defects in hESC-derived cerebral organoids.

## Results

### Characterization of RTT hESC-derived ASTs

To elucidate the function of MECP2 in the development of the human brain, we used both female and male hESC-derived cells in 2D and 3D cultures. The WIBR3^MGT^ is a female reporter line described in An et al., with GFP or tdTomato independently inserted in frame after *MECP2* exon 3 of each X allele, resulting in early termination and gene disruption^11^. For the second line, we generated an early stop mutation with our male hESC line to model one of the most common mutations in RTT (**Extended Data Fig. 1a,b**)^12^. Using CRISPR- Cas9 gene editing, linker eGFP was inserted after arginine 168 in WIBR1 hESC, resulting in early termination and a truncated LOF protein. These lines correspond to isogenic controls WIBR3 and WIBR1, respectively^13^. The following studies utilized the two control and two RTT hESC lines for analysis. Differentiation of hESC lines to mature ASTs was confirmed with expression of AST markers NDRG2, GFAP and EAAT2 (**Extended Data Fig. 1c**). We used a multivariate assessment to measure AST reactivity. First, ASTs were seeded at low density to examine cytoskeletal reorganization (**Fig. 1a, Extended Data Fig. 1c**). Measuring dimensions of the cell cytoskeleton with βeta-actin, we found that RTT AST have a smaller surface area compared to control (CTR) ASTs (**Fig. 1a, Extended Data Fig. 1d**). Quantifying mitochondria labeled by mitochondrial import receptor subunit TOM20, we found a lower abundance of mitochondria with fewer branches and branch end points in RTT ASTs compared to CTR (**Extended Data Fig. 1e**). To examine potential differences in AST reactivity and metabolism, we tested a panel of compiled reactivity markers by quantitative PCR (qPCR)^14^ (**Fig. 1b**). Overall, we found a trend that RTT ASTs have shifted gene expression in both negative and positive directions. Similar to reactive changes observed in disease pathology^14^, RTT ASTs have increased levels of transcripts involved in metabolism (*TSPO*) and intermediate filaments (*GFAP* and *VIM*). Mitochondrial encoded NADH dehydrogenase 5 (*MT-ND5*) was the most significantly upregulated gene (**Fig. 1b**). Measuring AST proliferation, RTT ASTs show a small but significant increase in proliferation at 72 hours after seeding (**Extended Data Fig. 1f**). Together, these data suggest RTT AST undergo morphological, molecular, and functional remodeling indicative of AST reactivity in response to *MECP2 LOF*.

**Fig. 1:**
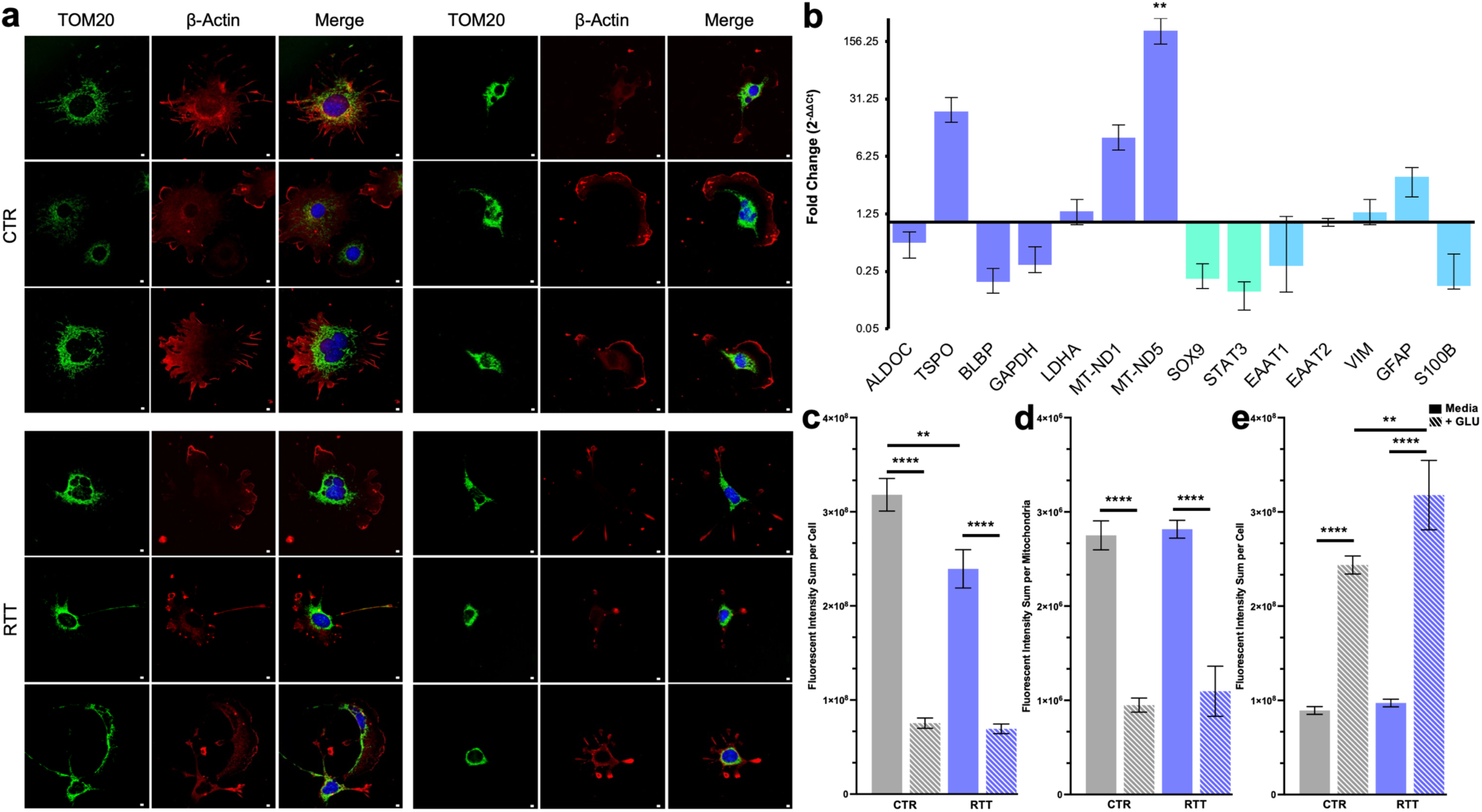
RTT AST characterization and function. **a)** Representative images of CTR (top panel) and RTT (bottom panel) ASTs seeded at a low density. Mitochondria (TOM20), Actin cytoskeleton (β-Actin). Merge with DAPI. Scale bar 5 µm. **b)** Quantitative PCR panel of AST reactive markers^14^ (*ALDOC, TSPO, BLBP, SOX9, STAT3, EAAT1, EAAT2, VIM, GFAP, S100β*). Marker colors: metabolism and mitochondrial (purple), transcription factors (green), and AST (blue) markers in RTT compared to CTR. Transcripts normalized to 18S. CTR n = 8, RTT n = 8, performed in triplicate per value. Statistical analysis Two-Way ANOVA Multiple Comparisons Bonferroni **p ≤ 0.01. Error bars Standard Error of Measure (SEM). **c – e)** Quantification of MitoTracker or MitoSOX in the absence (media) or presence (+ GLU) 200 µM GLU for 24 hrs. **c)** MitoTracker fluorescence per cell. Images for quantification Media CTR n = 32, RTT = 32, + GLU CTR n = 32, + GLU RTT n = 17. **d)** MitoTracker fluorescence per mitochondria. Media CTR n = 32, RTT = 32, + GLU CTR n = 31, + GLU RTT n = 16. **e)** MitoSOX fluorescence per cell. Media CTR n = 32, RTT = 32, + GLU CTR n = 17, RTT n = 17. Statistical analysis Two-Way ANOVA Multiple Comparisons Bonferroni **p ≤ 0.01, ****p ≤ 0.0001. Error bars SEM.

An essential function of an AST is GLU clearance at the excitatory synaptic cleft. Okabe et al. have previously identified diminished GLU clearance in cultured mouse Mecp2-deficient ASTs^4^, and Sun et al. found decreased GLU clearance in MECP2 R270X hESC-derives ASTs^10^. We found diminished GLU clearance in human RTT ASTs compared to CTR (**Extended Data Fig. 1g**), aligning with decreased expression of excitatory amino acid transporter *EAAT1* (**Fig. 1b**). We went on to examine the effect of adding a high concentration (200 µM, **Extended Data Fig. 1g**) of GLU over 24 hrs. Besides being an important neurotransmitter, GLU is an essential energy source in the brain that feeds into the mitochondrial TCA cycle, or is amidated to glutamine and released from ASTs for neuronal uptake^15^. We labeled mitochondria with MitoTracker or labeled superoxide, a byproduct of oxidative phosphorylation (OxPhos), with MitoSOX to quantify mitochondrial mass and activity, respectively (**Extended Data Fig. 1h**). We identified a significant decrease in MitoTracker per cell in RTT ASTs, confirming the observed decrease in mitochondrial mass from **Extended Data Fig. 1d** (**Fig. 1c**). There was no significant difference in MitoTracker intensity per mitochondria between RTT and CTR ASTs (**Fig. 1d**), which indicates similar labeling of MitoTracker. GLU treatment significantly decreased MitoTracker quantification in both CTR and RTT, suggesting changes to mitochondrial membrane potential after prolonged GLU treatment (**Fig. 1c,d**). Interestingly, we found no significant difference in MitoSOX fluorescence comparing RTT to CTR ASTs, indicating similar levels of superoxide production despite decreased abundance of mitochondria in RTT (**Fig. 1e**). GLU treatment significantly increased MitoSOX in both CTR and RTT, as predicted with increased mitochondrial function, but to a higher degree in RTT ASTs compared to CTR. These findings indicate decreased mitochondrial mass and similar levels of reactive oxygen species (ROS) in RTT ASTs compared to CTR.

### Defective mitochondrial function and altered energy metabolism in RTT ASTs

Mitochondria are critical organelles for metabolism in eukaryotes, generating both energy and precursors for biosynthesis of the essential building blocks of the cells. Beyond their metabolic roles, mitochondria are increasingly recognized as important signaling organelles, and mitochondrial dysfunction has been associated with major diseases including cancer, diabetes, and neurodegeneration^16, 17^. RTT has many similarities to mitochondrial diseases, and reports have indicated altered mitochondrial membrane potential and respiration in muscle and the brain^18^. From the critical changes in mitochondrial abundance, ROS and mitochondrial gene expression in RTT AST compared to CTR (**Fig. 1, Extended Data Fig. 1**), we therefore examined mitochondrial and glycolytic function by Seahorse assay. Overall, oxygen consumption rate (OCR) was significantly lower, with the levels in RTT ASTs remaining below those of CTR (**Fig. 2a,b**). Treatment with ATP synthase inhibitor oligomycin did not significantly change OCR, suggesting little ATP-linked respiration and production. The most significant deficit was identified in maximal respiration, where RTT ASTs show little change in OCR compared to basal levels (**Fig. 2b**). This suggests RTT ASTs operate at near maximal capacity and resort to glycolysis to compensate for mitochondrial deficits. We confirmed this by a glycolysis stress test, and found significantly increased glycolysis and glycolytic capacity by increased extracellular acidification rates (ECAR) compared to CTR ASTs (**Fig. 2c,d**). Our lab previously showed a small reduction in basal oxygen consumption and maximal respiration rate in hESC-derived RTT cortical neurons, but no change in proton leak, adenosine triphosphate (ATP) production, nor glycolysis^19^. Altered bioenergetics in RTT ASTs was further exemplified by decreased total ATP levels (**Fig. 2e**). Together, RTT AST show significant defects in mitochondrial function which is compensated with glycolysis, yet glycolysis alone cannot compensate for the diminished ATP levels that are overall lower compared to CTR ASTs.

**Fig. 2:**
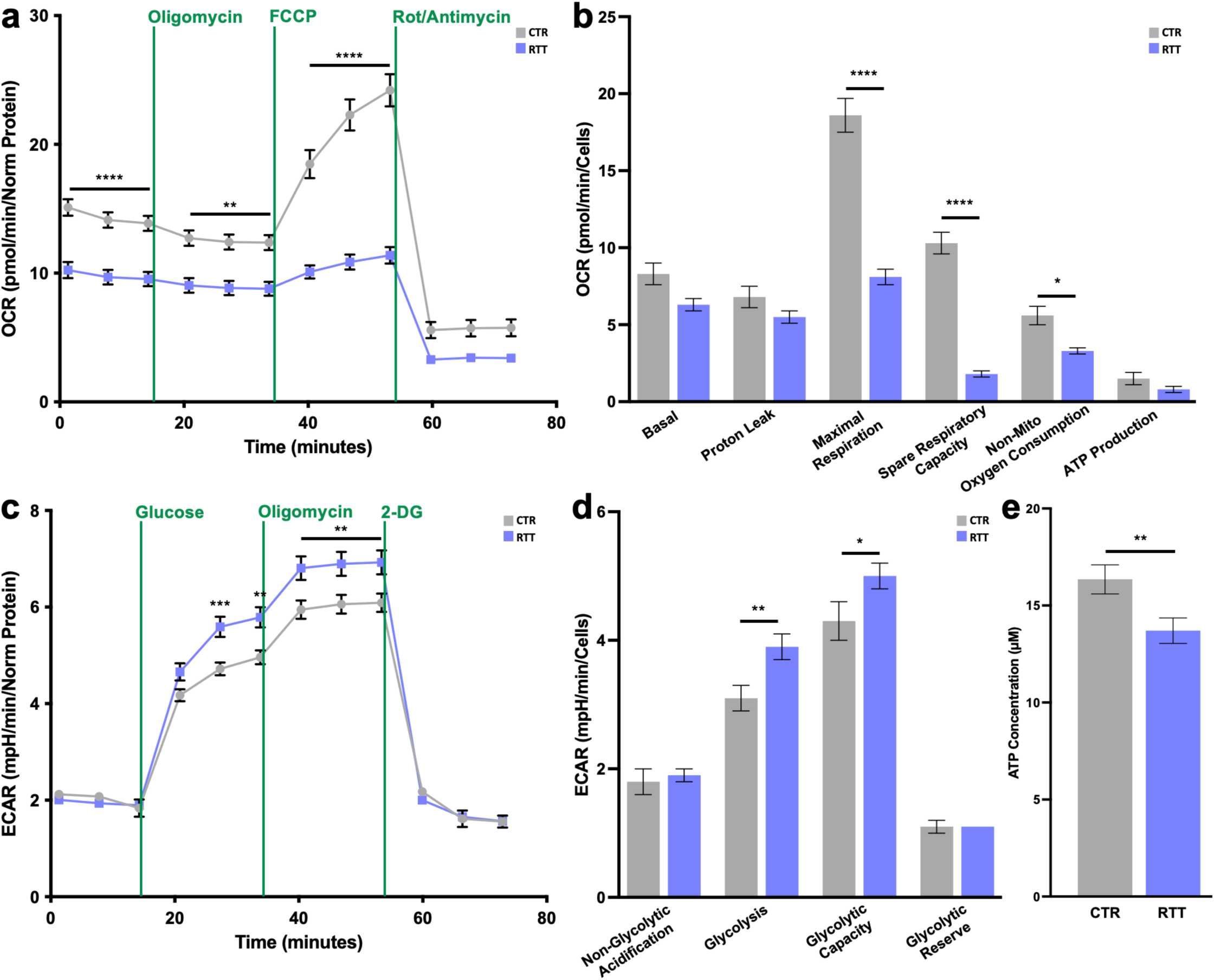
Dysfunctional mitochondrial respiration in RTT AST is compensated with increased glycolysis. **a)** Mitochondria stress test of CTR and RTT ASTs. To evaluate mitochondrial function, cells were injected with oligomycin (inhibit complex V), carbonyl cyanide-4-(trifluoromethoxy)phenylhydrazone (FCCP, collapses proton gradient), and rotenone/antimycin A (inhibits complexes I and III). CTR n = 46, RTT n = 46, Statistical analysis Two-Way ANOVA Multiple Comparisons Bonferroni **p ≤ 0.01, ****p ≤ 0.0001. Error bars SEM. **b)** Quantification of the basal oxygen consumption rate (OCR), proton leak, maximal OCR, spare capacity, Non-mitochondrial OCR, and ATP-linked OCR, in CTR and RTT AST **(a)**. Statistical analysis Two-Way ANOVA Multiple Comparisons Bonferroni *p ≤ 0.05, ****p ≤ 0.0001. Error bars SEM. **c)** Glycolysis stress test. Glycolysis was increased with glucose and oligomycin injection, followed by blocking of glycolysis with competitive glucose analog 2-DG. CTR n = 46, RTT n = 46. Statistical analysis Two-Way ANOVA Multiple Comparisons Bonferroni **p ≤ 0.01, ***p ≤ 0.001. Error bars SEM. **d)** Quantification of the extracellular acidification rate (ECAR) Non-glycolytic acidification, glycolysis, glycolytic capacity and reserve, in CTR and RTT ASTs **(c)**. Error bars SEM. Statistical analysis Two-Way ANOVA Multiple Comparisons Bonferroni *p ≤ 0.05, **p ≤ 0.01. **e)** Total ATP levels from CTR and RTT ASTs. CTR n = 72, RTT n = 72. Statistical analysis unpaired T-Test. Error bars SEM.

### Integration of proteomic and metabolomic profiles uncovers mechanism of mitochondrial dysfunction

Defining the molecular changes that arise from MECP2 LOF and downstream pathways that are affected is extremely difficult due to the multifaceted role of this protein. Recently, studies have identified biomarkers that correlate with metabolic dysfunction in autism spectrum disorders (ASD)^20, 21^. What is not well characterized in the ASD field are the metabolic changes at the cellular level in the brain, specifically how altered metabolites impact cell function and maturation. We examined potential biochemical and molecular differences between RTT and CTR ASTs with whole-cell metabolite and proteomic profiling. Whole-cell polar metabolomic analysis identified significant changes in metabolite abundances in glycerophospholipid metabolism (increased phosphocholine), glycolysis (increased lactate) and pentose phosphate pathway (decreased UDP-hexose) in RTT ASTs compared to CTR (**Extended Data Fig. 2a**). Interestingly, the abundance of numerous amino acids was significantly increased, including proline, arginine, methionine, threonine, and tryptophan (**Fig. 3a**). Through transamination or deamination reactions, amino acids can be converted into intermediates of TCA cycle. Performing whole cell proteomic analysis, we resolved 3451 proteins, and roughly half of the proteins (1465) showing significant abundance changes in both positive and negative directions in RTT compared to CTR (**Fig. 3b, Extended Data Fig. 2b**). Focusing on proteins involved in metabolism and mitochondrial function, we found the glycolysis/gluconeogenesis pathway significantly impacted, with many enzymes in the pathway increased including ALDOA and ENO (**Fig. 3c**). This agrees with the upregulation of glycolysis observed from the Seahorse assay (**Fig. 2c,d**). Many proteins that are localized in the mitochondria were altered (**Fig. 3d**), including proteins involved in the TCA cycle (**Fig. 3e**), and a large decrease in protein abundance for those involved in OxPhos, including Complex I (NUA4L, NDUFV1, NDUV2, NDUFS2, NDUFS4, NDUFS8, NDUFS3, NDUF7, NDUFS1) and ATP synthase Complex V (ATPG, ATPD, ATPB, ATPA, ATPF2) (**Fig. 3f**). We also investigated amino acid metabolism and degradation pathways after observing the increased amino acids from whole cell metabolomics in RTT AST. We found that proteins significantly altered in arginine, proline, cysteine, methionine, glycine, serine, threonine, tryptophan metabolism, and lysine degradation (**Extended Data Fig. 2c-g**), were shifted down, and few changes lactate metabolism (**Extended Data Fig. 2h**).

**Fig. 3:**
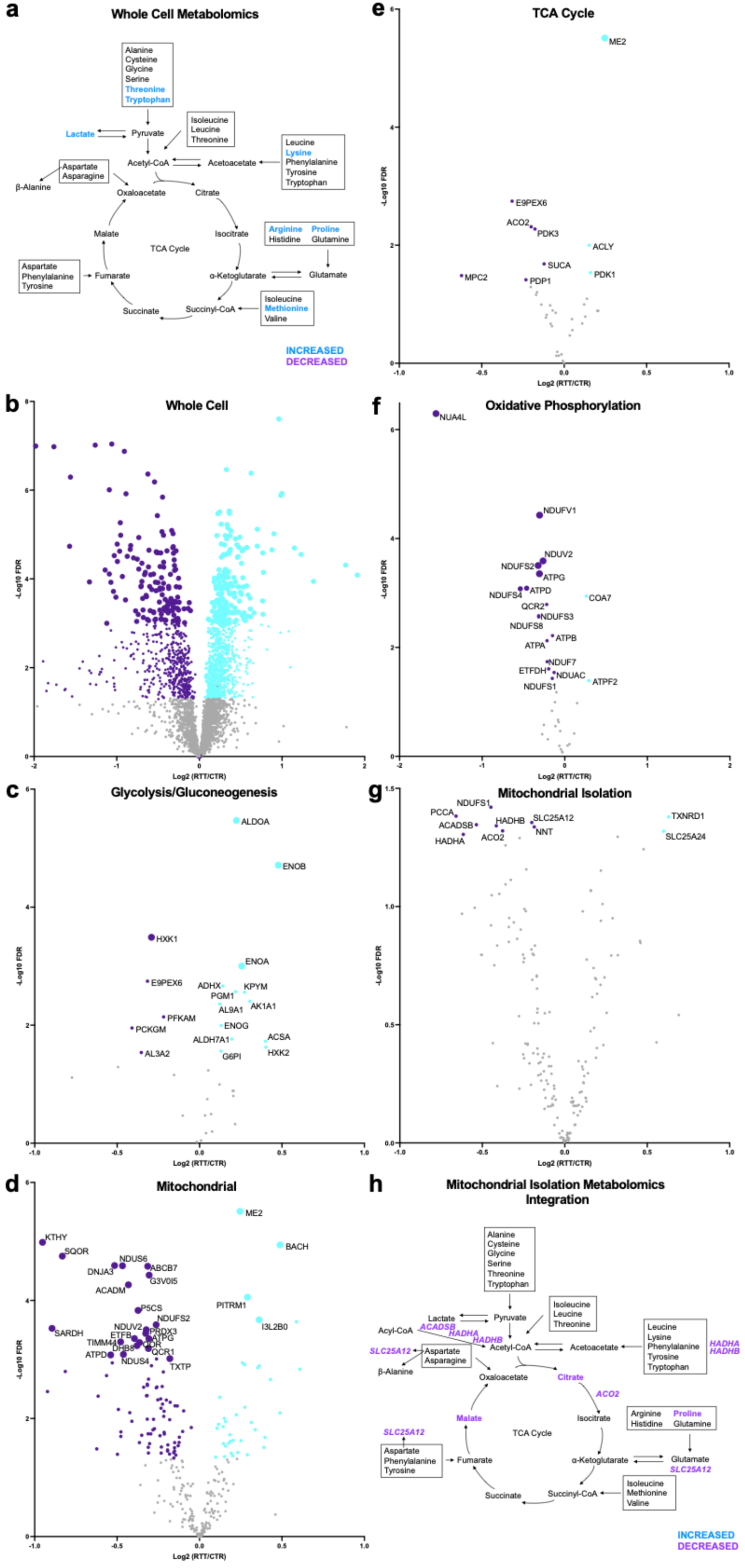
Integration of metabolomics and proteomics analyses indicates molecular changes tied to mitochondrial dysfunction in RTT AST a) Whole cell AST metabolomics summary. TCA cycle with metabolite changes of RTT relative to CTR (increased in blue, unchanged in black). b-f) Volcano plots of whole cell AST proteomic analysis. Total protein changes of RTT compared to CTR (b), and protein changes in specific metabolic pathways for glycolysis/gluconeogenesis (c), mitochondrial localized proteins (only adjusted ***p ≤ 0.001 labeled) (d), TCA Cycle (e), and oxidative phosphorylation (f). Increased proteins (light blue), decreased proteins (purple) in RTT compared to CTR. Large dot adjusted ***p ≤ 0.001, small dot adjusted *p ≤ 0.05. CTR n = 6, RTT n = 6. g) Volcano plot of isolated mitochondria from ASTs proteomic analysis. We resolved 174 proteins. Total protein changes of RTT compared to CTR. Increased proteins (light blue), decreased proteins (purple) in RTT compared to CTR. Small dot adjusted *p ≤ 0.05. CTR n = 6, RTT n = 6. h) Isolated mitochondria from ASTs proteomics and metabolomics integration summary. TCA cycle with metabolite changes of RTT relative to CTR (decreased in purple, unchanged in black). Metabolites in normal font, proteins in italicized font.

To examine the molecular mechanism tied to energy dysfunction in RTT ASTs, we probed the proteome and metabolome of isolated mitochondria. Fractionation of mitochondria was confirmed by western blot analysis, with mitochondrial import proteins VDAC and TOM20 found exclusively in the mitochondrial fraction and not cytosolic (**Extended Data Fig. 3a**). As we observed a significantly lower mitochondrial mass in RTT AST, we normalized the isolated mitochondria proteomics by loading the same amount of total protein.

We indeed found significantly altered protein abundances similar to whole cell proteomics, including decreased NDUFS1 of Complex I in OxPhos, and HADHA and HADHB in amino acid metabolism (**Fig. 3g**). We then probed the metabolome of the isolated mitochondria. We resolved similar metabolites compared to previously published human MITObolome^22^, and discovered diminished abundance of metabolites involved in amino acid metabolism (phosphocholine and proline), cofactor/redox metabolism (1-methylnicotinamide and nicotinamide), nucleotide metabolism (S-adenosylmethionine), pentose phosphate pathway (6-phosphogluconate and UDP-hexose) and the TCA Cycle (citrate and malate) (**Extended Data Fig. 3b,c**). We then integrated the mitochondrial proteomic and metabolomic analyses, and found the proteins correlate to perturbations in metabolites, converging around the TCA cycle and ETC (**Fig. 3h**). Within the TCA cycle, Aconitase 2 (ACO2) converts citrate to isocitrate. Blocking ACO2 in colorectal cancer cells caused a reduction in TCA cycle intermediates, suppression of mitochondrial OxPhos, increased glycolysis, and elevated citrate flux for fatty acid and lipid synthesis^23^. An additional connection is the decrease of nicotinamide and 1-methylnicotinate metabolites and decrease of NADH:ubiquinone oxidoreductase core subunit S1 (NDUFS1) and nicotinamide nucleotide transhydrogenase (NNT) proteins, supporting redox imbalance disruption of the electron transport chain (ETC) (**Extended Data Fig. 3b**,**c**, **Fig. 3g)**. Further, a group found that NNT knockdown significantly reduced ACO2 activity in a non-small cell lung cancer line, leading to a reduced capacity to drive ETC flux^24^. Together, we defined biochemical evidence tied to mitochondrial energetic dysfunction after MECP2 LOF in ASTs.

### RTT ASTs rely on amino acid catabolism as compensatory mechanism for diminished energetics

To test how RTT ASTs adapt to environmental changes, we probed the response to metabolic shift. Testing three conditions that will alter mitochondrial function, we exchanged glucose with either galactose (GAL), found to stimulate OxPhos, or with lipid cocktail (LIP), found to stimulate the TCA cycle, or stimulated glycolysis by removing pyruvate from the media (-PYR). We performed whole cell metabolomics on ASTs after incubation with conditions for 24 hours (**Extended Data Fig. 4).** Overall, we found minimal changes to metabolites in CTR AST. We found mitochondrial stimulation in CTR ASTs depletes levels of lactate and few amino acids, while glycolysis stimulation in CTR ASTs depletes ATP levels compared to basal media (**Fig. 4a, Extended Data Fig. 4**). When we examined the RTT ASTs, we found a large shift in metabolites in all conditions compared to basal media. We observed similar changes compared to the shift in CTR AST metabolites, however, there was a significant decrease in a staggering number of metabolites overall. This includes metabolites in the pentose phosphate pathway (UDP-hexose GAL, LIP) and pyrimidine metabolism (beta-alanine, GAL, LIP, -PYR) and TCA Cycle (citrate) that were not observed in CTR ASTs. Of particular interest, we identified decreased abundance of metabolites in all amino acid metabolism pathways (**Fig. 4a, Extended Data Fig. 4**). These findings indicate that RTT ASTs deplete levels of almost every amino acid that feeds into the TCA cycle, along with glutamate and lactate after energy demands are shifted.

**Fig. 4:**
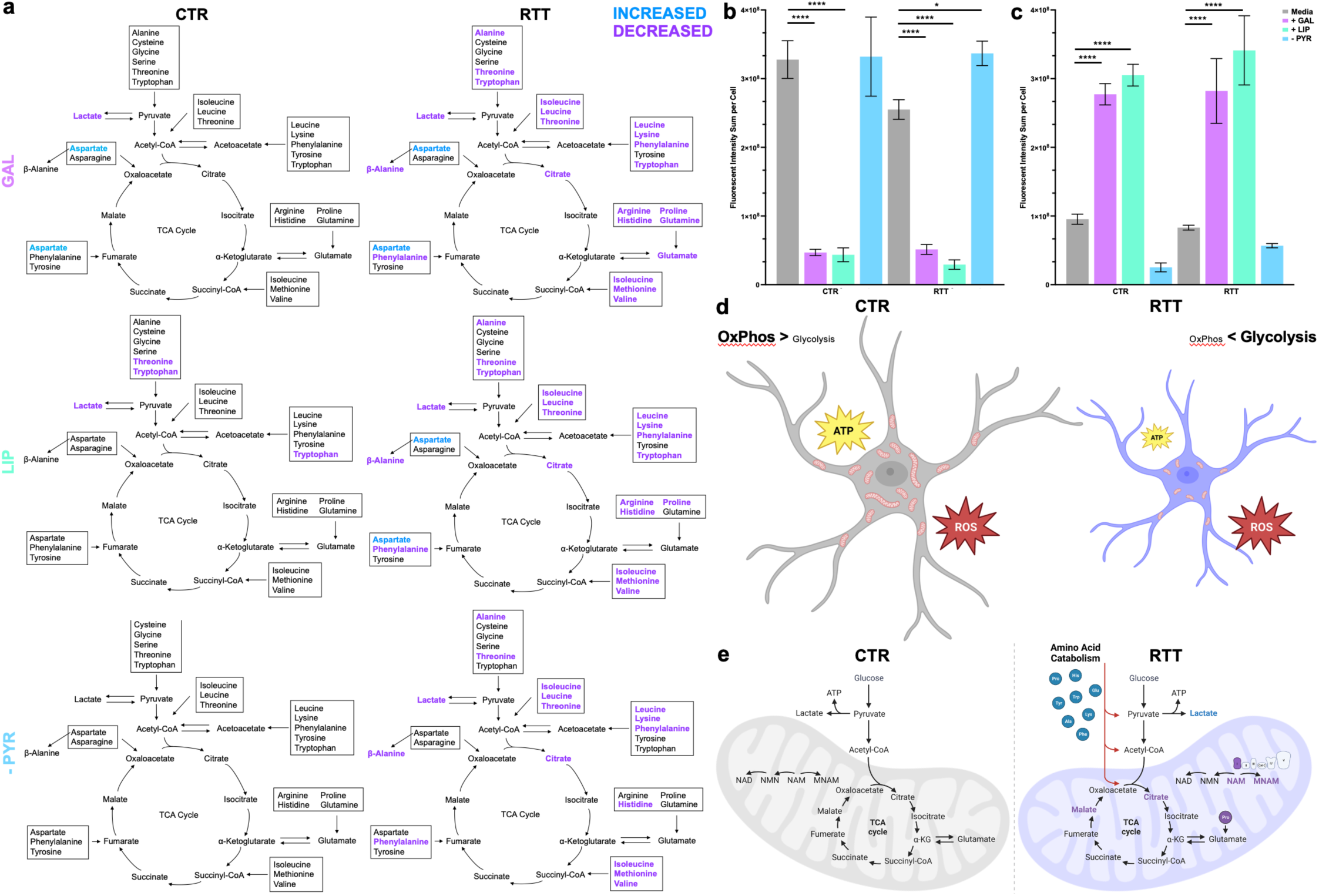
RTT AST compensate for altered energy demands by catabolizing amino acids. **a)** Whole cell AST metabolomics summary. TCA cycle with metabolite changes relative to glucose media control (increased in blue, decreased in purple) in CTR (left) and RTT (right). Media conditions on left side. GAL – galactose exchanged for glucose, LIP - lipid cocktail exchanges for glucose, - PYR - removal of sodium pyruvate. **b-c)** Quantification of MitoTracker or MitoSOX after incubation with media conditions for 24 hrs. **b)** MitoTracker fluorescence per cell. Images for quantification CTR Media n = 16, CTR GAL n = 16, CTR LIP n = 7, CTR –PYR n = 15, RTT Media n = 32, RTT GAL n = 15, RTT LIP n = 15, RTT –PYR n = 32. **c)** MitoSOX fluorescence per cell. CTR Media n = 32, CTR GAL n = 31, CTR LIP n = 15, CTR –PYR n = 16, RTT Media n = 16, RTT GAL n = 15, RTT LIP n = 14, RTT –PYR n = 31. Statistical analysis Two-Way ANOVA Multiple Comparisons Bonferroni *p ≤ 0.05, ****p ≤ 0.0001. Error bars SEM. **d)** Model summarizing Figs. 1 and 2, **Extended Data** Fig. 1. RTT AST indicate altered reactivity, decreased mitochondrial mass, similar levels of ROS, and decreased ATP production. With dysfunctional mitochondria, they compensate with increased glycolysis over oxidative phosphorylation (OxPhos). Created with BioRender.com. **e)** Model summarizing **Figs. 2-4**, **Extended Data Fig. 2-4.** Mitochondria indicate diminished levels of metabolites in TCA cycle and in redox balance (proteomic analysis). Whole cell ASTs have increased abundance of amino acids and lactate, which is depleted when energy demands are increased. NAD nicotinamide adenine dinucleotide, NMN nicotinamide mononucleotide, NAM nicotinamide, MNAM methyl-nicotinamide. Created with BioRender.com.

Next, we examined mitochondrial response to metabolic shift by confocal microscopy. After changes to media for 24 hours, as indicated above, we labeled mitochondria with MitoTracker or labeled superoxide with MitoSOX (**Extended Data Fig. 1h**). As shown in **Fig. 1c**, we observed decreased mitochondrial mass in RTT compared to CTR under basal media (**Fig. 4b**). In CTR and RTT ASTs, mitochondrial stimulation (GAL, LIP) decreased MitoTracker labeled mitochondria, while glycolysis stimulation (-PYR) had no impact on CTR, yet increased MitoTracker labeled mitochondria in RTT ASTs (**Fig. 4b**). Decreased mitochondrial function may improve mitochondrial membrane potential in RTT AST, explaining the increased MitoTracker fluorescence.

Examining superoxide abundance, we observed a significant increase in MitoSOX in mitochondrial stimulating conditions in both CTR and RTT AST (**Fig. 4c**). From the increased ROS production, we propose that RTT ASTs can increase mitochondrial function when stressed by increasing catabolism of amino acids. We summarize our findings on how MECP2 LOF dramatically impacts RTT AST energetics and amino acid metabolism in RTT AST in **Fig. 4d,e**.

### Dysfunctional mitochondria from RTT ASTs are transferred to cortical neurons and augment LFP activity

Identifying mitochondrial dysfunction in RTT ASTs potentially has an even greater impact on brain development as ASTs provide significant energy support to neurons. ASTs can donate their mitochondria to neurons, and vice-versa, through tunneling nanotubes, extracellular vesicles and gap junctions, an essential response to neuronal excitability and injury^25^. To examine if MECP2 LOF impacts the ability of ASTs to donate their mitochondria to neurons, and the impact of MECP2 LOF on the ability of neurons to take up extracellular mitochondria, we labeled mitochondria in ASTs with MitoTracker and then seeded cells onto 2 month cultured cortical neurons. After 48 hours, cultures were fixed and stained to identify colocalization of MitoTracker with neurons. We identified labeled mitochondria in CTR and RTT neurons from CTR and RTT ASTs, indicating that RTT ASTs can donate their mitochondria and RTT neurons can uptake mitochondria (**Fig. 5a)**. Inversely, we examined the ability of neurons to donate their mitochondria to ASTs. Labelling neuronal mitochondria with MitoTracker, we seeded ASTs onto neurons and examined colocalization of MitoTracker with ASTs. Similarly, we identified labeled mitochondria in CTR and RTT ASTs from CTR and RTT neurons (**Fig. 5b**). We found similar correlation of MitoTracker colocalization in the recipient cells (**Extended Data Fig. 5a**).

**Fig. 5:**
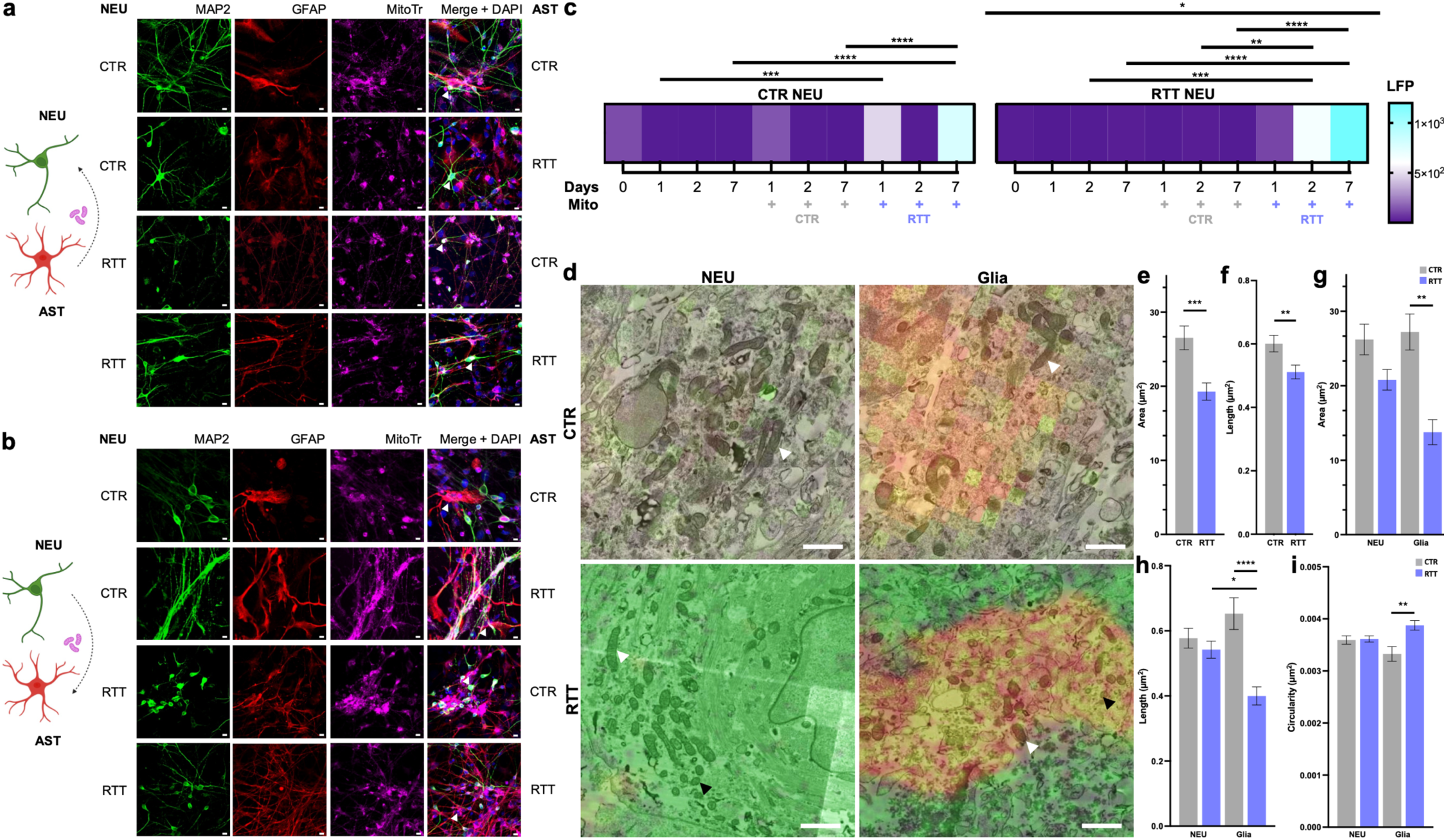
Dysfunctional mitochondria from RTT AST can transfer to neurons and cause hyperexcitability. **a)** Representative images of transfer of mitochondria labeled in ASTs to hESC-derived cortical neurons (NEU). AST mitochondria labeled with MitoTracker, followed by seeding AST onto NEU cultures. Left column, genotype of NEU. Right column, genotype of AST. MitoTracker positive mitochondria identified in NEU (white arrowhead). NEU (MAP2), AST (GFAP), MitoTracker (MitoTr), Merge with DAPI. Scale bar 5 µm. Schematic created with BioRender.com. **b)** Representative images of transfer of mitochondria labeled in NEUs to ASTs. NEU mitochondria labeled with MitoTracker, followed by seeding AST onto NEU cultures. Left column, genotype of NEU. Right column, genotype of AST. MitoTracker positive mitochondria identified in AST (white arrowhead). NEU (MAP2), AST (GFAP), Mitotracker (MitoTr), Merge with DAPI. Scale bar 5 µm. Schematic created with BioRender.com **c)** Multielectrode Array (MEA) analysis of NEU cultures with AST mitochondria. Mitochondria were isolated from CTR and RTT AST and seeded onto NEU. Local field potential (LFP) activity was measured over 7 days. CTR and RTT LFP activity did not significantly differ. After addition of CTR mitochondria (+ CTR), LFP activity did not significantly differ. After addition of RTT mitochondria (+ RTT), LFP activity significantly increased in both CTR and RTT NEUs. CTR Day 0 n = 20, Day 1 n = 11, Day 2 n = 9, Day 7 n = 9, CTR + CTR mito Day 1 n = 5, Day 2 n = 4, Day 7 n = 5, CTR + RTT mito Day 1 n = 11, Day 2 n = 9, Day 7 n = 7, RTT Day 0 n = 28, Day 1 n = 9, Day 2 n = 10, Day 7 n = 10, RTT + CTR mito Day 1 n = 2, Day 2 n = 3, Day 7 n = 3, RTT + RTT mito Day 1 n = 14, Day 2 n = 14, Day 7 n = 14. Statistical analysis Two-Way ANOVA Multiple Comparisons Bonferroni with **Extended Data Fig. 6a.** *p ≤ 0.05, **p ≤ 0.01, ***p ≤ 0.001****p ≤ 0.0001. **d)** Representative images of Correlative Light Electron Microscopy (CLEM) on Day 100 matured CTR and RTT cerebral organoids. Left, mitochondria in NEU (green). Right, mitochondria in glia (AST/oligodendrocytes) (red). Healthy fused mitochondria (white arrowhead) and small, circular fission mitochondria (black arrowhead). Scale bar 1 µm. **e-i)** Quantification of mitochondria ultrastructure morphology by CLEM. Significant change in total area **(e)** and length **(f)** between CTR and RTT organoids. Statistical analysis T-Test **p ≤ 0.01, ***p ≤ 0.001. Error Bars SEM. **(g-i)** Comparing mitochondria in NEU and glia, significant decrease observed in mitochondria from glia area **(g)** and length **(h)** in RTT compared to CTR, decrease in mitochondria length between NEU and glia in RTT **(h)**, and increased circularity of mitochondria in glia in RTT compared to CTR **(i)**. Statistical analysis One-Way ANOVA Tukey *p ≤ 0.05, **p ≤ 0.01. Total CTR n = 251, total RTT n = 430, NEU CTR n = 172, NEU RTT n = 337, Glia CTR n = 79, Glia RTT n = 93. Error bars SEM.

To examine the consequence of mitochondrial transfer from RTT ASTs to neurons, we designed a mitochondrial transplantation assay. Utilizing an updated method and different lines than our previous report^19^, we first examined phenotypic differences between CTR and RTT hESC-derived cortical neurons. We recapitulated the same phenotype that we previously reported, with decreased soma size and less intersections by sholl analysis in RTT compared to CTR neurons^19^ (**Extended Data Fig. 5b,c**). Isolated mitochondria from CTR and RTT ASTs were then seeded onto CTR and RTT cortical neuron cultures.

MitoTracker labeled mitochondria were robustly taken up by both CTR and RTT neurons after 48 hours (**Extended Data Fig. 5d**). Importantly, isolated mitochondria exhibited good integrity and purity, as indicative of retention of MitoTracker dye. We then seeded isolated mitochondria from CTR and RTT ASTs onto CTR and RTT cortical neurons on a multielectrode array plate, where local field potential (LFP) activity was measured over 7 days (**Fig. 5c, Extended Data Fig. 6a**). LFP activity was similar between CTR and RTT neurons over the course of the experiment in the absence of exogenous mitochondria. After addition of exogenous mitochondria isolated from CTR ASTs onto CTR and RTT neurons, we observed relatively no change in activity compared to neurons without mitochondria (**Fig. 5c**). However, adding exogenous mitochondria isolated from RTT ASTs onto CTR and RTT neurons resulted in significant changes to LFP activity. This activity was visualized as roughly 6-to-8-fold increase in LFPs for CTR and RTT neurons relative to neurons without exogenous mitochondria. Additionally, the frequency of LFPs in RTT compared to CTR neurons seeded with RTT mitochondria at Day 7 was significantly increased, indicating RTT mitochondria increase RTT neuronal LFP activity to a higher degree than observed in CTR neurons. Examining neuronal ATP levels after 48 hours post mitochondrial seeding, we found the CTR and RTT mitochondria have no impact on total ATP levels, indicating that increased LFP activity is not due to changes in ATP abundance (**Extended Data Fig. 6b**). These findings illustrate that dysfunctional mitochondria from RTT ASTs can be transferred to neurons, impacting activity in a disease context.

**Fig 6:**
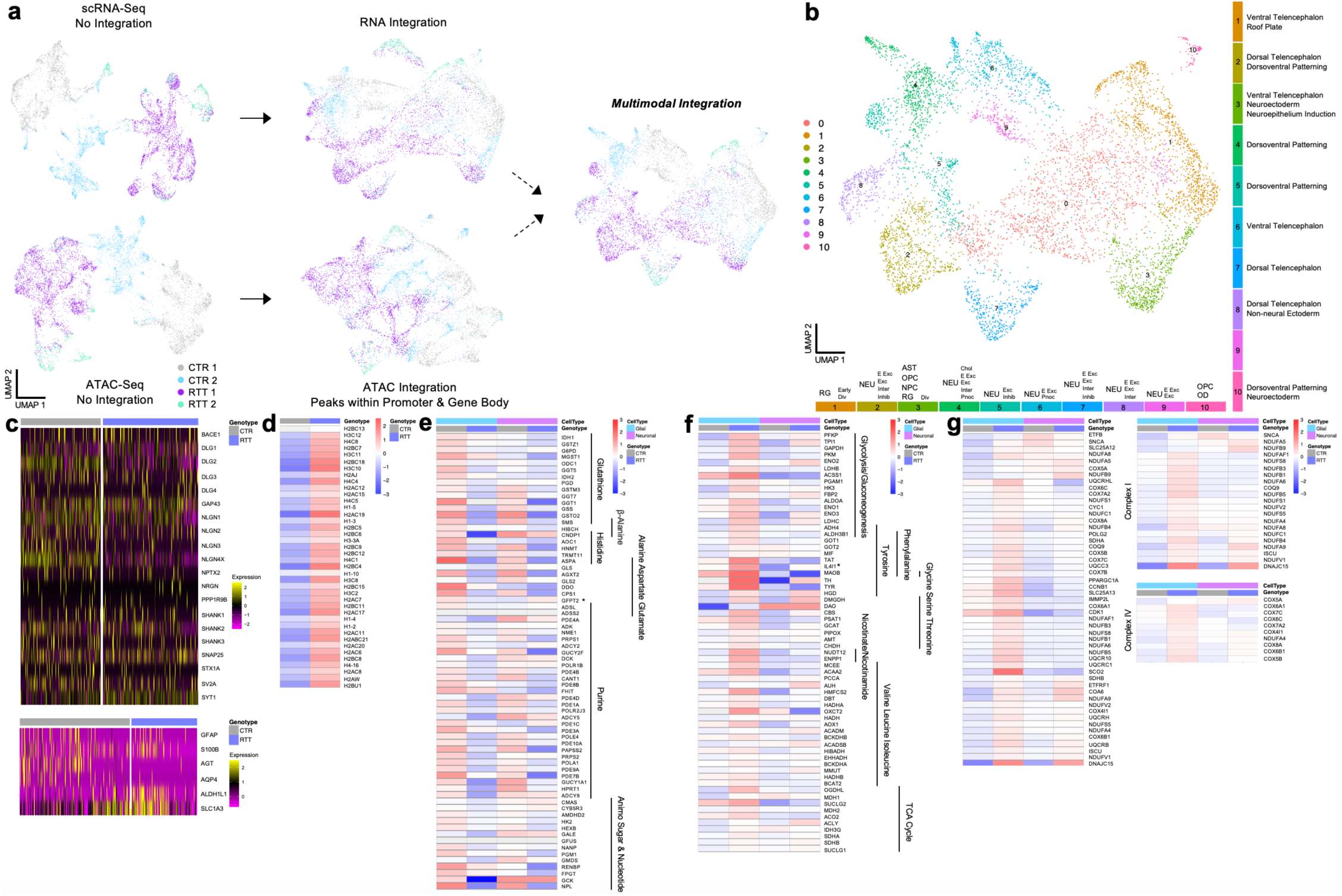
Multiomics analysis on RTT cerebral organoids reveals distinct transcriptional and epigenetic changes that differ in neuronal and glia populations. a) Uniform manifold approximation projection (UMAP) plot embedding of integrated multiomics. Two CTR (grey and blue) and two RTT (purple and green) organoids matured to day 100 were utilized for the experiment. Single nucleus (sn) RNA-seq and snATAC-seq UMAP prior to integration indicates strongest pattern is the difference in expression from the genotype. After integration, snRNA-seq and snATAC-seq modalities were combined for multimodal integration. b) UMAP plot colored by cluster. Due to heterogeneity of cluster 0, this was further subclustered and defined in Extended Data Fig. 7b,c. Bottom legend cell identities: Radial Glia (RG), RG Early, RG Division (Div), Astrocyte (AST), Oligodendrocyte Precursor Cell (OPC), Neural Progenitor Cell (NPC), Oligodendrocyte (OD), Neuron (NEU), Early Excitatory NEU (E Exc), Excitatory NEU (Exc), Interneuron (Inter), Inhibitory NEU (Inhib), Cholinergic NEU (Chol), Pnoc NEU (Pnoc positive). Right legend early brain anatomy. c) Heat map of neuron and AST markers. Each column represents one cell. Clusters containing only neurons (0,2,4-9) (top) for markers of activity and mature synapses, and only glia and progenitor cells (1,3,10) (bottom) for AST markers. d-g) Heatmaps showing gene expression, CTR (left), RTT (right). d) Genes enriched on the structural constituent of chromatin gene set. Normalized enrichment score (NES) 2.65 and FDR 0, p-value 0 on the GSEA ranking genes by differential expression. e) Combined Kyoto Encyclopedia of Genes and Genomes (KEGG) Metabolism GSEA sets enriched in glia CTR population compared to glia RTT population. Genes within sets (identified on right) were combined. *GFPT2 – also in Amino Sugar and Nucleotide Metabolism. f) Combined KEGG Metabolism GSEA sets enriched in glia RTT population compared to glia CTR population. Genes within sets (identified on right) were combined. *IL4I1 – also in Valine Leucine Isoleucine Metabolism. g) GSEA sets enriched in glia RTT population compared to glia CTR population. Genes enriched on the (Left) Respiratory ETC – NES 2.07, FDR 0.008, p-value = 0. (Right) Mitochondrial ETC NADH to Ubiquinone (Complex I). NES 1.91, FDR 0.043, p-value = 0.0001. Mitochondrial ETC Cytochrome C to Oxygen (Complex IV) NES 1.58, FDR 0.356, p-value = 0.038.

### Mitochondrial morphology defects in RTT hESC derived cerebral organoids

To study the impact of MECP2 LOF on human brain development in a more physiologically relevant system, we generated cerebral organoids from the RTT lines and isogenic CTRs. We matured unguided neural organoids to 100 days in culture, a timepoint that matches neuronal modules at prenatal stages *in vivo*^26^. In order to both visualize mitochondria and differentiate cell types, we performed correlative light electron microscopy (CLEM) on the female RTT and CTR pair (W3 and W3^MGT^). We fixed and imaged organoids after incubation with live cell dyes to label neurons green and ASTs/oligodendrocytes red. After imaging by scanning transmission electron microscopy, we overlayed the fluorescent images onto the electron microscopy images and quantified mitochondrial dynamics (**Fig. 5d**). Overall, we found that mitochondria from RTT organoids were significantly smaller compared to mitochondria from CTR organoids. Total area and length were decreased in RTT compared to CTR (**Fig. 5e,f, Extended Data Fig. 6c-e**). Separating the mitochondria based on cell type, we found a stronger phenotype in RTT glia (**Fig. 5g,h, Extended Data Fig. 6f,g**). In addition to total area and length differences, we found increased circularity in mitochondria from RTT glia compared to CTR, suggesting a shift towards either mitochondrial fission or reduced fusion, as mitochondria are not branched and forming the same networks as CTR (**Fig. 5i**). Measuring the total area of cristae per mitochondria, there were no statistically significant differences comparing RTT to CTR, but there was a trend for decreased cristae area in glia compared to neurons (**Extended Data Fig. 9e,g**). Together, this analysis indicates that RTT glia developed in a model for human brain development have altered mitochondrial morphology that corroborates observed mitochondrial dysfunction from RTT AST monoculture. We suggest that the smaller mitochondria observed in RTT neurons may originate from RTT AST.

### Developmental delay and altered metabolic gene enrichment in RTT cerebral organoids

MECP2 was the first methyl-binding protein discovered^27^, and since then the role of MECP2 in epigenetic regulation has been studied extensively^28–32^. However, we do not fully understand the impact of *MECP2* mutations on gene expression and chromatin landscapes in the human brain, and how MECP2 functions as the brain develops. To address these areas, we used a multiomics approach on the female RTT and CTR isogenic pair organoids at Day 100. We performed single-nucleus RNA-seq (snRNA-seq) plus Assay for Transposase-Accessible Chromatin (ATAC)-seq (snATAC-seq) on barcoded DNA and mRNA from isolated nuclei. To define the exact composition of every cell in a single organoid, we performed the multiomics technique on two individual RTT organoids compared to two individual CTR organoids. Prior to integration, there is clear separation between CTR and RTT organoids when examining spatial expression of snRNA-seq and snATAC-seq uniform manifold approximation projection (UMAP)s. (**Fig. 6a**). We can deduce that while organoids can be heterogeneous, the *MECP2* mutation is the driving factor behind the variability between cells. Next, we integrated the snRNA-seq and snATAC-seq data, followed by multimodal integration of both datasets that was utilized for all analyses (**Fig. 6a**). We identified 11 cell clusters using the smart local moving SLM algorithm^33^ on the weighted combination of gene expression and ATAC-seq peak signal multimodal neighbors as implemented in Seurat v4.1.1.9001^34^ (**Fig. 6b, Extended Data Fig. 7a**). Cluster 0 was further subclustered into 3 Clusters (**Extended Data Fig. 7b,c**). To both infer identities of each cluster and compare cell type specification across datasets, we utilized two human organoid datasets from Camp et al.^35^ and Fleck et al.^36^, a human cortex dataset from Nowakowski et al.^37^, and the BrainSpan Atlas of the Developing Human Brain^38^ via Cell-type Specific Expression Analysis tool developed by Dougherty *et al*.^39^ as reference atlases to find consensus across different developmental timepoints and high confidence in cell identity annotation. We inferred cell type identity, cell maturation stage and anatomical semblance to the developing human cortex. We identified glial populations in clusters 1, 3, and 10, with early and dividing radial glia in cluster 1, ASTs, oligodendrocyte precursor cells, neural progenitor cells and dividing radial glia in cluster 3, and a separate population of oligodendrocyte precursor cells and oligodendrocytes in cluster 10. Clusters 2, 4-9, and subclusters 0-2 contained a heterogeneous neuron population which included early excitatory, excitatory, intern, inhibitory, cholinergic, and Pnoc+ neurons (**Fig. 6b, Extended Data Fig. 7b**). We also mapped anatomical features of the human developing cortex by cluster (**Fig. 6b, Extended Data Fig. 7b**). There is clear separation of dorsal telencephalon (clusters 2, 7-8, and subcluster 0) and ventral telencephalon (clusters 1, 3, 6). Additional features included presence of roof plate (cluster 1 and subcluster 2) dorsoventral patterning (clusters 2, 4-5, 10, and subcluster 1), neuroepithelium induction (cluster 3), neuroectoderm (clusters 3 and 10), and non-neural ectoderm (cluster 8). Anatomical classification highlights the diversity of cells based on cluster cell type identify. Examining the proportion of cells per cluster based on genotype, we found that the absence of cluster 10 from RTT organoids indicates RTT organoids did not develop mature oligodendrocytes by this timepoint (**Fig. 6b**, **Extended Data Fig. 7b,d**).

Next, we focused on transcriptional expression differences between CTR and RTT organoids in the neuron and glia/progenitor (referenced as glia) populations by combining annotated clusters. Overall, we identified differing cell type proportions in organoid composition, with an increased abundance of neurons and decreased abundance of glia in RTT (80% neuron and 20% glia) compared to CTR (66% neuron and 34% glia) (**Supplemental Table 1**). We note there was a similar total number of CTR (3636) and RTT (3648) cells.

Common markers for neuron maturity, including genes encoding for synaptic and activity markers, had lower expression in RTT organoids compared to CTR (**Fig. 6c**). Common AST markers are expressed at lower levels in RTT organoids compared to CTR, and also lack expression of markers *AGT* and *AQP4,* indicating not all subpopulations of AST are present at this timepoint in RTT organoids^40^ (**Fig. 6c**). These results indicate an increased abundance of immature neurons and decreased abundance of glia with distinct glial subtypes, including oligodendrocytes (**Extended Data Fig. 7d**), not developed in RTT organoids compared to CTR. Together, our RTT organoids share many phenotypes as RTT ASTs and epitomize the characteristic developmental delay observed as clinical features in RTT patients^41^, organoids^42, 43^ and mouse models^44, 45^.

We found both significantly up- and down-regulated genes in RTT organoids (**Extended Data Fig. 8a,b**). Our group has previously reported a reduction in overall gene expression after *MECP2* mutation in cortical neurons using spike in normalization^19^. We observed a decrease in global gene expression as previously reported^19^ but still detected several significantly upregulated genes with large increases in gene expression in RTT organoids (**Extended Data Fig. 8c)**. Differential gene expression trends were similar in the neuron and glial populations (**Extended Data Fig. 8d,e**). Examining gene set enrichment analysis (GSEA) from total differential gene expression using the C5 ontology gene sets, we identified Structural Constituent of Chromatin as the most significantly enriched, containing both core and linker histones in RTT compared to CTR (**Fig. 6d**). We then used GSEA to investigate enrichment of gene sets in genes differentially expressed between RTT and CTR in the neuron and glial populations separately using the C5 ontology and the Kyoto Encyclopedia of Genes and Genomes (KEGG) subset of canonical pathways gene sets. Genes were selected if they were part of the core enrichment on the GSEA for KEGG genes sets related to metabolism. Examining KEGG gene sets enriched in glial population of CTR organoids, we found increased genes for metabolism of glutathione, purine, amino sugar and nucleotides, and amino acids (**Fig. 6e**). Examining KEGG gene sets enriched in neuron population of CTR organoids, we found similar overlap but also increased galactose, pyruvate, and nitrogen, and differences in amino acid metabolism (**Extended Data Fig. 9a**). We then examined KEGG and gene ontology biological process gene sets enriched in glial population of RTT organoids, and found increased genes for glycolysis, nicotinate/nicotinamide, TCA cycle and amino acids (**Fig. 6f**), respiratory ETC, and specific sets to ETC Complex I and IV (**Fig. 6g**). Importantly, these sets were not enriched in the RTT neuron population. We found different amino acid metabolism and degradation, and metabolism of pyrimidines enriched in neuron population of RTT compared to CTR (**Extended Data Fig. 9b**). These findings directly correlate with our metabolomic, proteomic, and Seahorse assay RTT AST data (**Figs. 2-4**), indicating glial specific enrichment of metabolic and mitochondrial pathways in RTT organoids that were also perturbed in RTT AST monocultures.

To trace transcriptional changes from the multiomics study to translational outcome and examine morphology of the cells, we stained male and female hESC-derived organoids with common neuronal and AST markers. We observed a large abundance of neurons in RTT organoids compared to CTR (**Extended Data Fig. 9c)**. Examining pre- and post-synaptic markers, we identified lower expression in RTT compared to CTR, indicating much of these NeuN positive neurons are immature and lack canonical pre- and post-synaptic markers as indicated in **Fig. 6c** (**Extended Data Fig. 9c**). We identified a similar abundance of S100β and GFAP positive ASTs between RTT and CTR, however there is a significant difference in AST morphology.

ASTs in CTR organoids display extended processes that are completely absent in RTT organoids (**Extended Data Fig. 9c**). We suggest these ASTs are immature or have altered reactivity, and together these results recapitulate the developmental delay that is characteristic in RTT. Our results also agree with previously published findings in human cerebral organoid RTT models at earlier stages that identified decreased neurite growth, neurogenesis and defective neural progenitor migration^42, 46–48^.

### Global changes in RTT cerebral organoid chromatin accessibility

We investigated the DNA regions that MECP2 most significantly modifies, either directly or indirectly, in the developing brain. With our multiomics analysis, we found ATAC-seq peaks correlated well with gene expression when comparing differentially expressed genes in glia and neurons in CTR and RTT organoids (**Extended Data Fig. 10a**). We found more peaks that have increased chromatin accessibility in RTT compared to CTR organoids (**Extended Data Fig. 10b,c**). Linking peaks at 3 kb or less to proximal genes, we examined potential changes in differentially accessible peak locations relative to the gene body in CTR and RTT glial and neuronal populations. Overall, small differences were observed between the glia and neuron differentially accessible peak locations when comparing immediate promoter and intragenic regions in CTR and RTT (**Extended Data Fig. 10d,e**). The most significant phenotype was observed between regions more accessible in CTR and more accessible in RTT organoids. We found a difference in the distribution of differentially accessible peaks at the immediate promoter (52% and 44% peaks more accessible in RTT compared to 29% and 21% peaks more accessible in CTR) in glia and neurons, respectively. Conversely, peaks with higher accessibility in CTR organoids were ∼2.3 and ∼2.7x more likely to be located in distal intergenic regions for neurons and glia, respectively (**Extended Data Fig. 10d,e**). Together, we found that with loss of MECP2 function, the epigenetic landscape in developing human organoids is shifted to more accessible chromatin with differences in the genomic locations of accessibility.

## Discussion

Much research throughout the past decade has identified strong correlations between brain pathologies and glial dysfunction^49–51^. We provide multiple lines of evidence for mitochondrial dysregulation and dysfunction in RTT ASTs, including increased *MT-ND5* expression, lower abundance of mitochondria that produce similar levels of superoxide, broad defects in mitochondrial respiration, and diminished levels of total ATP. We identified a compensatory mechanism for mitochondrial dysfunction via increased glycolysis and catabolism of amino acids when energy demands are increased in RTT ASTs. Dysfunctional mitochondria from RTT ASTs can be transferred to CTR and RTT hESC-derived cortical neurons, directly impacting neuronal activity. We found that in a 3D culture to model the developing brain, mitochondria display morphology deficits that suggest diminished activity in hESC-derived cerebral organoids. Several papers have concluded that RTT ASTs secrete factors that impair neuron function^6–8, 52, 53^. We propose that dysfunctional mitochondria released from RTT ASTs may also be a contributing factor to the discoveries from these papers. For instance, mitochondria can readily be transferred between cells by gap junction internalization^54^, and previous findings by Maezawa et al. in a *Mecp2^-/+^* deficiency mouse model that RTT can spread from ASTs through connexin 43 (Cx43)- mediated gap junctions. When Cx43 was knocked down with siRNA, the spread of MeCP2 deficiency was significantly reduced^52^. Our findings illuminate how healthy mitochondria in ASTs are critical to the developing brain, and open a new therapeutic avenue for rescue of RTT AST dysfunction by pharmacological intervention or mitochondrial transplantation^55^.

Overall, our study widely agrees with mitochondrial defects in RTT mouse models, including total decreased ATP levels^56–58^ and increased ROS^59–62^. However, our study suggests significant differences between human and mouse ASTs. Cultured ASTs from an *Mecp2* null mouse indicated increased ROS, yet differed with increased mitochondrial mass and no change in size nor response to mitochondrial uncoupling with FCCP^63^. Recent studies have indicated the most significant species-specific differences between human and mouse ASTs lie in mitochondrial resting state, physiology, and increased susceptibility of oxidative stress^64^. We also shed new light on how MECP2 function greatly differs between neurons and glia in the developing brain. We used a multiomics approach on cerebral organoids to measure mRNA and chromatin accessibility from the same nuclei, an approach that has not been previously reported on RTT tissue or cells. Through multimodal integration, this technique provided a comprehensive understanding of the molecular changes due to MECP2 LOF in developing cerebral organoids. By cross-comparing 4 separate references to define clusters, we hold high confidence in the composition and anatomy of the organoids. We found MECP2 LOF not only impacts neuronal development and maturation, but also the abundance and subtypes of glia.

This highlights the importance of defining MECP2 function based on cell type, molecular context, and species. We propose upregulation of genes involved in metabolism and mitochondrial function in the glial population is a compensatory mechanism (**Figs. 1b, 6f,g**). With largely diminished proteins surrounding oxidative phosphorylation, TCA cycle, ETC, and amino acid metabolism (**Fig. 3b-f, Extended Data Fig. 2**), we propose a feedback loop where RTT glia increase gene expression to compensate for mitochondrial dysfunction and altered metabolism.

Many groups, including ours, have proposed that MECP2 functions as both a transcriptional repressor and activator due to direct and indirect interactions with RNA, DNA, and protein. Our previous studies have identified decreased total gene expression after MECP2 mutation in whole cell analysis of cortical neurons^19, 65^. We see this pattern as well, with a downward shift of gene expression in RTT compared to CTR when the normalization is not based on RNA-seq sequencing depth (**Extended Data Fig. 8c**). However, we identified a higher proportion of genes and accessible peaks that were significantly increased in RTT compared to CTR organoids. We also found significant enrichment of genes encoding for histones in RTT organoids compared to CTR (**Fig 6d**) indicating that disrupting MECP2 has a global epigenetic impact by upregulation of histone complexes. These findings provide novel insight into MECP2 function in the developing human brain.

## Methods

### hESC Cell Lines and Maintenance

hESCs were maintained feeder-free in mTeSR Plus media (STEMCELL Tech 100-0276) on Matrigel coated plates (Corning 354234). Passaging of hESC was performed with either ReLeSR (STEMCELL Tech 100-0484) or manual passaging. Mycoplasma contamination was routinely checked for all cells in culture. Target vector for this knock-in strategy utilized pUC19 plasmid (Addgene) with homology directed repair (Plasmid pCL186_pUC19_HDR_hMECP2-GFP-R168X). Sanger sequencing confirmed that a linker-eGFP transgene is inserted after MECP2-R168 in the Exon 4 position to result in a early truncation (**Extended Data Fig. 1a**). Western blot confirmed truncated MECP2 with absence of C-terminus (**Extended Data Fig. 1b**). WIBR3^MGT^ was a generous gift from Dr. Thor Theunissen, Washington University in St. Louis. Unless noted, all experiments used both CTR and RTT lines for analysis.

### Stem Cell Differentiation and Maintenance

#### Astrocytes

Stem cells were transitioned to Glial Progenitor Medium, modified from Muffat et al. 2018^66^, containing Neurobasal (Life Technologies 21103049), DMEM/F12/HEPES (Thermo Fisher Scientific 12400024), BSA-v (7.5%) (Gibco 15260-037), GlutaMAX (Life Technologies 35050061), Sodium Pyruvate (Life Technologies 11360070), N-2 (Life Technologies 17502048), 20 ng/mL CNTF (Peprotech AF-450-13-500UG), Penicillin Streptomycin. ASTs were passaged with Accutase (StemCell Technologies 07920), and maturity was confirmed with expression of markers in **Extended Data Fig. 1a**, roughly 150 days from 1^st^ media change.

#### Astrocyte Metabolite Media

AST base media (**Fig. 4**). Neurobasal -D-glucose -sodium pyruvate (Life Technologies A2477501), BSA-v, GlutaMAX, N-2, CNTF, Penicillin Streptomycin. Conditions: *Media* - add 25 mM D-Glucose (Sigma G7021) and sodium pyruvate. *Media + GAL* - add 25 mM D-Galactose (Sigma G0625) and sodium pyruvate. *Media + LIP* - add 1/40 lipid cocktail (gibco 11905-031) and sodium pyruvate. *Media - PYR* - add 25 mM D-Glucose and same volume of Neurobasal -D-glucose -sodium pyruvate to exchange sodium pyruvate amount.

#### Cortical Neurons

Neurons were generated according to protocol from Fernandopulle et al. 2018^67^. Briefly, px330-AAVS1-DOX- NGN2-mcherry and px330-guideAAVS1-GFP plasmids were used for gene targeting of NGN2 knock-in at the AAVS1 locus. Lipofectamine Stem Transfection Reagent (Life Technologies STEM00008) and Opti-MEM Reduced Serum (Life Technologies 31985070) were used for transfection of plasmids into hESC lines. Cells were selected with puromycin, diluted to single cell and screened by expression of mcherry. Multiple clones were tested for each genotype. After Accutase treatment, 1.5 x 10^6^ cells were seeded in a 6-well plate. After 3 days of Induction Media with Doxycycline, cells were treated with Accutase and seeded onto Matrigel coated plates for final neuron differentiation. Neurons were maintained in BrainPhys Maturation Media, modified from Fernandopulle et al. 2018^67^ to include Culture One supplement (ThermoFisher Scientific, A3320201) to maintain a homogeneous population of cortical neurons. Neurons were matured for at least 2 months for analysis. Axion Biosystems 48-well plates (M768-tMEA-48B), 1 x 10^5^ cells. 96-well imaging plates (ThermoFisher Scientific 165305), 1 x 10^4^ cells.

#### Cerebral Organoids

Organoids were constructed with STEMdiff Cerebral Organoid kit (STEMCELL Tech 08570). After one month from EB seeding, organoids were slowly transferred to maturation media containing Neurobasal, Gem21 (GeminiBio 400-160) Glutamax, Penicillin Streptomycin. At 5 weeks, organoids were cut at 350 microns to prevent necrotic tissue in the center. Cutting buffer contained HBSS - Ca^2+^ (Life Technologies 14175095) HEPES (Gibco 15630-080) Glucose and Penicillin Streptomycin. Cutting of organoids continued routinely every four weeks until organoid diameter maintained constant size was no larger than 500 microns. Roughly, organoids were cut 4 times previous to Day 100 time point for assays with a McIlwain Tissue Chopper at 350 microns. Organoids were maintained in hypoxic chamber at 37°C on an orbital shaker at 100 RPM.

### Immunocytochemistry, Immunohistochemistry and Microscopy

#### Astrocytes

For **Fig. 1** and **Extended Data Fig. 1,** ASTs were seeded at a low density (100/well) in 96-well imaging plate. After 24 hours, ASTs were washed once in phosphate buffered saline solution (PBS) and fixed in fresh 4% paraformaldehyde (PFA) in PBS for 1 hour at room temperature. ASTs were washed multiple times and permeabilized with PBST (1 × PBS solution with 0.1% Triton X-100), then blocked in PBST with 10% BSA for 1 hour at room temperature. Primary antibodies (GFAP anti-mouse Dako M0761, NDRG2 abcam ab174850- 100µL, EAAT2 anti-mouse Santa Cruz Biotechnology sc-365634, Beta-Actin anti-mouse Proteintech 60008-1- IG, TOMM20 anti-rabbit abcam ab186734-100µL) were added at 1:500 in PBST with 5% BSA overnight at 4°C. After washing, secondary antibodies (488 anti-mouse Jackson 715-545-151, 594 anti-rabbit Jackson 711- 585-152) and DAPI (Life Technologies D1306) were added at 1:1000 in PBST with 5% BSA overnight at 4°C. After washing, ProLong Diamond Antifade Mountant (Invitrogen P36961) was added to wells prior to imaging. Images were captured on a Zeiss LSM700 confocal microscope and processed with Zen software and ImageJ/Fiji. For imaging-based quantification, unless otherwise specified, at least 12 representative images per genotype/condition were quantified and data were plotted as mean ± SEM with Excel or Graphpad Prism.

#### MitoTracker and MitoSOX Assay and Analysis

MitoTracker deep red (ThermoFisher Scientific, M22426) was diluted to 500 nM in AST media and incubated on ASTs for 1 hour in chamber. MitoSOX Red (ThermoFisher Scientific, M36008) was diluted to 5 µM in HBSS + Ca^2+^ and incubated on ASTs for 10 min in chamber. Cells were washed multiple times before fixation. Images were captured on a Zeiss LSM980 airyscan microscope, Z-stacked and processed with Zen software and ImageJ/Fiji. For imaging-based quantification, unless otherwise specified, at least 12 representative images per genotype/condition were quantified and data were plotted as mean ± SEM with Excel or Graphpad Prism.

### Mitochondrial Transfer between Astrocytes and Neurons

Cortical neurons were matured for at least 2 months prior to imaging. ASTs or neurons were incubated with MitoTracker for 1 hour, followed by washing and passaging of ASTs onto neurons (1 x 10^4^ cells per well).

Fixation and antibody protocol is described above. Primary antibodies (MAP2 anti-chicken Encor Biotechnology CPCA-MAP2, GFAP anti-rabbit), secondary antibodies (488 anti-chicken Jackson 703-545-155, 594 anti-rabbit) plus DAPI. For isolated mitochondrial transplantation from ASTs to neurons, isolated mitochondria (see protocol below) were incubated with MitoTracker for 10 min rocking at room temperature.

After 48 hours, neurons were incubated with MitoSOX for 10 minutes, then washed multiple times in PBS and fixed with 4% PFA. Primary antibody MAP2 anti-chicken, secondary antibody 488 anti-chicken plus DAPI. Images were captured on a Zeiss LSM700 confocal microscope and processed with Zen software and ImageJ/Fiji. For imaging-based quantification, unless otherwise specified, at least 12 representative images per genotype/condition were quantified and data were plotted as mean ± SEM with Excel or Graphpad Prism.

#### Organoids

At Day 100, organoids were washed in PBS and fixed in 4% PFA in PBS. Organoids were preserved and cleared using SHIELD (Lifecanvas technologies C-PCK-250), then permeabilized in PBST and antibodies were added as above. Primary antibodies (NeuN anti-mouse EMD Millipore MAB377, GFAP anti-mouse, PSD-95 anti-rabbit abcam ab18258, S100B anti-rabbit Dako Z0311, SYT-1 anti-goat LSBio LS-B5899), secondary antibodies (488 anti-mouse, 594 anti-rabbit, 680 anti-goat Jackson 705-625-147) plus DAPI. Organoids were incubated in EasyIndex optical clearing solution for few hours and mounted on slides in same solution. Images were captured on a Zeiss LSM700 confocal microscope, Z-stacked and processed with Zen software and ImageJ/Fiji. At least 4 organoids were imaged per condition/genotype.

### qRT-qPCR

Whole cell ASTs: 5 x 10^5^ cells pelleted and snap frozen. RNA was extracted by RNeasy Plus Micro Kit (Qiagen 74034) and cDNA was synthesized by qScript cDNA Supermix (Quantabio 95048-100). qPCR prepared with Fast SYBR Green Master Mix (appliedbiosystem 2023-08-31) and run with QuantStudio 6.

### qPCR

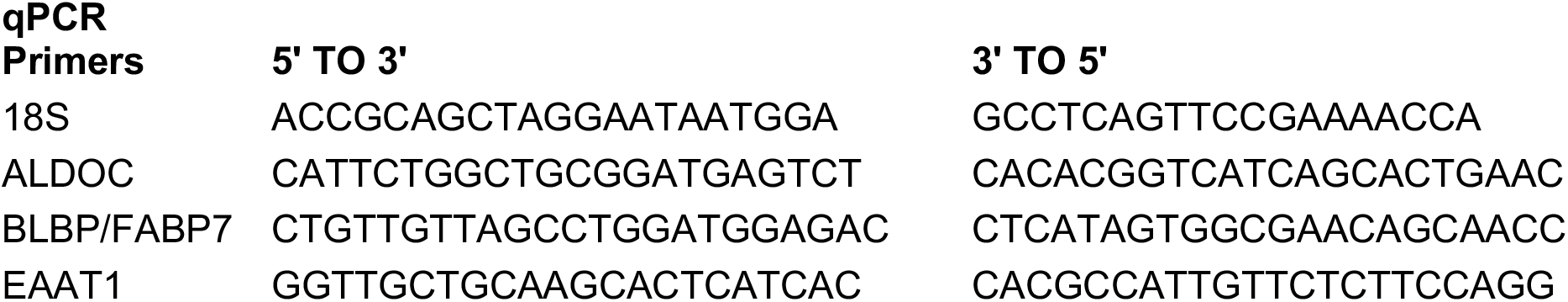
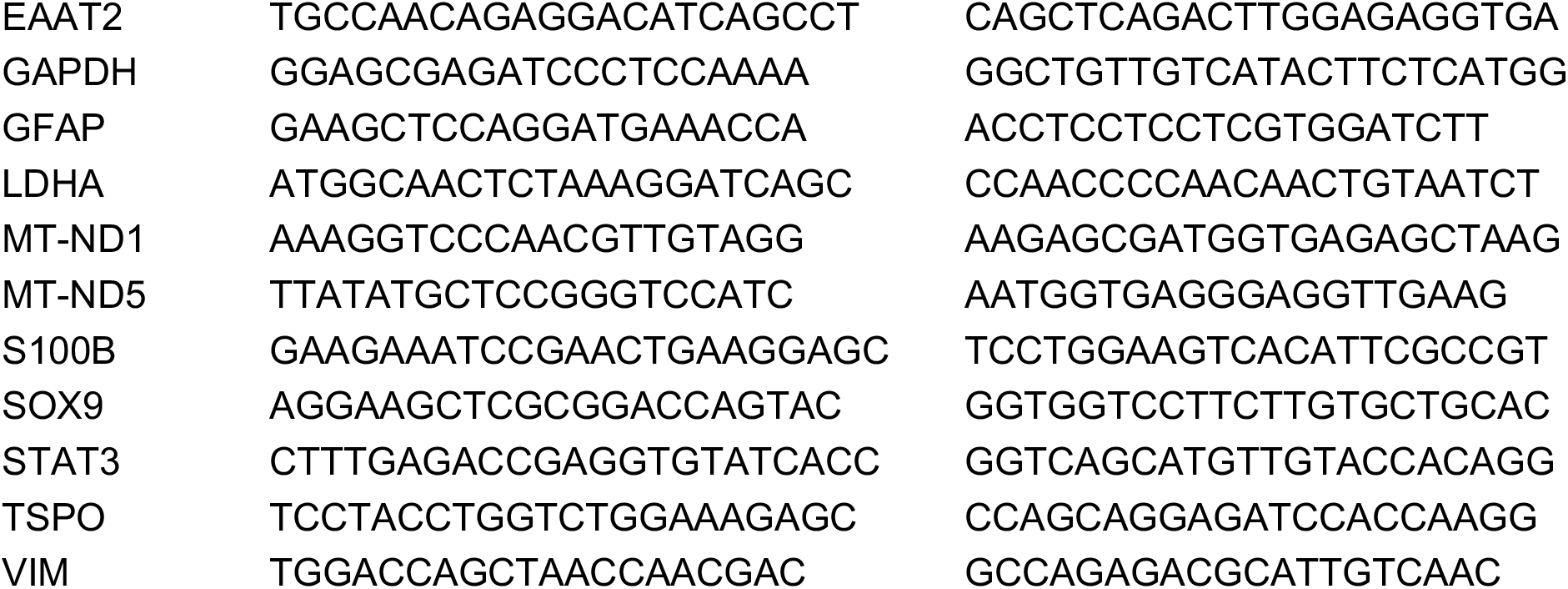

### Cell Analysis Proliferation

ASTs were seeded at 1 x 10^4^ cells in Matrigel coated 24-well plates. At 24, 48, and 72 hours, cells were dissociated with Accutase and counted.

### Image Analysis

Tom20 Analysis - Tom20 labeled mitochondria were analyzed with Fiji Image J plugin Mitochondria Analyzer for 2D analysis. Beta-Actin Analysis - Individual ASTs were outlined with the region-of-interest freehand tool in Fiji Image J.

### Glutamate Assay

ASTs plated at 1 x 10^5^ in 96-well plate. After 48 hours, cells were treated with 200 µM GLU for 4 hours. Glutamate assay was performed according to protocol with Sigma Glutamate Assay Kit (MAK004).

### ATP Assay

ASTs plated at 1 x 10^5^ in 96-well plate. After 48 hours, ATP assay was performed according to protocol with abcam ATP Bioluminescence Assay Kit (Ab113849).

### Proteomics

#### Protein extraction, TMT-labelling, solid phase extraction, and peptide fractionation

Isolated ASTs or mitochondria samples were lyophilized and resuspended in 23 µL S-trap lysis buffer (5% (w/v) SDS, 50 mM TEAB pH 8.5, 2 mM MgCl^2^, 10 mM TCEP, 0.025 U/µL Benzonase nuclease), vortexed, and incubated for 10 min at 55°C. The resulting lysate was centrifuged for 10 min at 20,000 x g and 4 °C and the supernatants were transferred to fresh Eppendorf tubes. Each sample was mixed with 5 µL 500 mM 2- iodoacetamide and the reactions were alkylated at room temperature for 30 min. After that, the protein extracts were acidified with 2.5 µL 27.5 % (w/v) phosphoric acid and mixed with 165 µL S-trap binding buffer (100 mM TEAB pH 7.55 in 90 % (v/v) ice-cold methanol). All liquid was applied to S-trap Micro columns (ProtiFi CO2- micro) and the protein suspensions were extracted via centrifugation for 1 min at 4000 x g. The columns were washed six times with 150 µL S-trap binding buffer and then the columns were placed into fresh Low Protein Binding Microcentrifuge Tubes (Thermo Fisher Scientific 90410). Trapped proteins were cleaved by adding 1

µg trypsin/LysC mix dissolved in 30 µL 50 mM TEAB pH 8.5 and incubating the digests over night at 37°C in a humified incubator. Tryptic peptides were eluted by centrifuging the columns for 1 min at 4000 x g followed by three washes with 40 µL 50 mM TEAB pH 8.5, 40 µL 0.2% (v/v) formic acid in water, and 40 µL 50% acetonitrile in water, respectively. The collected peptides were lyophilized and resuspended in 20 µL 100 mM TEAB pH 8.5. Peptide concentrations were determined using the Pierce Quantitative Fluorometric Peptide Assay (Thermo Fisher Scientific 23290) following the descriptions of the manufacturer. Then 2-8 µg peptide was dissolved in 20 µL 100 mM TEAB pH 8.5 and 100 µg TMTpro 16plex labels (Thermo Fisher Scientific A44522) dissolved in 5 µL anhydrous acetonitrile was added to each sample. The labelling reactions were incubated for 60 min at room temperature and subsequently quenched with 2 µL 5% (w/v) hydroxylamine for 30 min. From each labelled stock 10 µL were pooled in a fresh Low Protein Binding Microcentrifuge Tube, respectively. Pooled samples were lyophilized and resuspended in 100 µL 0.2% TFA followed by C18 solid phase extraction using stage tips following the descriptions by Rappsilber et al.^68^. Eluted peptides were lyophilized and reconstituted in 300 µL 0.2% TFA for fractionation using the Pierce High pH Reversed-Phase Peptide Fractionation Kit following the instructions provided by Thermo Fisher Scientific for TMT-labelled samples. The individual peptide fractions were lyophilized and reconstituted in 20 µL 0.2 % formic acid in water.

#### LC-MS

The collection of LC-MS/MS data was conducted using a Vanquish Neo nanoLC equipped with an Orbitrap Eclipse mass spectrometer, a FAIMS Pro Interface, and an Easy Spray ESI source (Thermo Fisher Scientific). An Acclaim PepMap (75μm x 2 cm) trap column together with a EasySpray ES803 column (75 µm x 50 mm, 100 Ȧ) (both Thermo Fisher Scientific) were employed for nanoLC with 3 µL peptide extract being injected.

Peptides were separated using 0.1% (v/v) formic acid in water (solvent A) and 0.1% (v/v) formic acid in 80% (v/v) acetonitrile (solvent B) as mobile phases with a 200 nL/min flow rate and a column temperature of 40°C. The column was conditioned for two minutes at 5% solvent B and then the linear gradient was gradually increased to up to 40% solvent B over 180 min. Remaining peptides bound to the C18 resin were then eluted for 16 min at 95% B. The ion source temperature was set to a temperature of 300°C in positive mode at 2100 V and ionized peptides were passed through by the FAIMS Pro unit at -45, -55, and -65 V. The mass spectra were collected with a resolution of 120,000 in DDA (data-dependent acquisition) mode at a mass range of m/z 400-1400, 300% automatic gain control (AGC) target and automatic injection time. The MS/MS fragmentation was performed by picking all precursor ions above 2.0 x 104 intensity at 0.7 Da m/z isolation windows and subjected to high energy collisional dissociation (HCD) using a normalized collision energy of 38. Each precursor ion already collected in MS2 mode was excluded from fragmentation for 60 s. Resulting MS2 product ions were analyzed with a 250% AGC target setting and 200 ms maximum ion accumulation time in the Orbitrap mass analyzer at 50,000 resolutions.

#### Data Analysis

Analysis of proteome data was conducted with the PEAKS Studio 10.6 software package. The data extracted from the .raw files was pre-processed using the following settings: merge scans (10 ppm retention time window; 10 ppm precursor m/z tolerance; precursor mass and charge states z = 2-8) followed by automatic centroiding, deisotoping, and deconvolution. A FASTA database for human proteins downloaded from Uniprot (UP000005640) or the MitoCarta 3.0 Database^69^ was used for protein identification under the following settings: parent mass error tolerance 10 ppm; fragment mass error tolerance 0.05 Da, retention time shift tolerance 5.0 min; semispecific trypsin enzyme specificity; variable modifications: carbamidomethylation (C), phosphorylation (STY), pyro-glu (Q), oxidation (M), deamidation (NQ); one non-specific cleavage specificity on one terminus, maximal two missed cleavages, maximal three variable posttranslational modifications per peptide. The decoy-fusion approach was utilized to determine the false discovery rates (FDRs) of peptide identifications. All peptide identifications that matched a false discovery rate of ≤ 1 % and a significance threshold of 20 (-10lgP) were classified as identified. The quantities of at least 2 of the most abundant peptide signals were utilized to compute the raw protein peak areas. The quantitative protein data was exported into the .csv format and analyzed as described previously^70^.

### Metabolomics

*Whole cell processing* - 1 x 10^6^ cells per well in 6-well plate, were washed with PBS. Cells were scraped in LC grade methanol and homogenized in Eppendorf tube containing water and LC grade chloroform with pestle mixer, followed by vortexing for 10 min at 4°C. Polar metabolites were separated by centrifuging top speed at 4°C. Samples were dried with a nitrogen dryer and stored at -80°C until all samples were ready, each sample is biological replicate.

*Isolated mitochondria processing* - 1.5 x 10^7^ cells were spun down for mitochondrial isolation (see protocol below). A small fraction of mitochondria per sample were lysed and tested for protein quantity by Bradford assay. Equal amounts of protein (5 µg) were added to individual eppendorf tubes for metabolite extraction as above. Each sample is technical replicate.

*Metabolite profiling* - conducted on a QExactive bench top orbitrap mass spectrometer equipped with an Ion Max source and a HESI II probe, which was coupled to a Dionex UltiMate 3000 HPLC system (Thermo Fisher Scientific). External mass calibration was performed using the standard calibration mixture every 7 days and an additional custom mass calibration was performed weekly alongside standard mass calibrations to calibrate the lower end of the spectrum (m/z 70-1050 positive mode and m/z 60-900 negative mode) using the standard calibration mixtures spiked with glycine (positive mode) and aspartate (negative mode). Typically, samples were reconstituted in 50 uL water and 2 uL were injected onto a SeQuant® ZIC®-pHILIC 150 x 2.1 mm analytical column equipped with a 2.1 x 20 mm guard column (both 5 mm particle size; Millipore-Sigma). Buffer A was 20 mM ammonium carbonate, 0.1% ammonium hydroxide; Buffer B was acetonitrile. The column oven and autosampler tray were held at 25°C and 4°C, respectively. The chromatographic gradient was run at a flow rate of 0.150 mL/min as follows: 0-20 min: linear gradient from 80-20% B; 20-20.5 min: linear gradient form 20- 80% B; 20.5-28 min: hold at 80% B. The mass spectrometer was operated in full-scan, polarity-switching mode, with the spray voltage set to 3.0 kV, the heated capillary held at 275°C, and the HESI probe held at 350°C. The sheath gas flow was set to 40 units, the auxiliary gas flow was set to 15 units, and the sweep gas flow was set to 1 unit. MS data acquisition was performed in a range of m/z = 70-1000, with the resolution set at 70,000, the AGC target at 1x10^6^, and the maximum injection time at 20 msec. All solvents, including water, were purchased from Fisher and were Optima LC/MS grade. Relative quantitation of polar metabolites was performed with TraceFinder™ 4.1 (Thermo Fisher Scientific) using a 5 ppm mass tolerance and referencing an in-house library of chemical standards. Data were filtered according to predetermined QC metrics: CV of pools <25%; R of linear dilution series <0.975. Metabolite enrichment analysis was performed with www.metaboanalyst.ca.

### Seahorse Analysis

ASTs were seeded onto Seahorse Cell Culture 96-well plate at 3 x 10^4^ cells per well. After 24-hours, Seahorse analysis was carried out according to Agilent protocol. Concentrations: Oligomycin 1.5 µM, FCCP 1 µM, Rot/Antimycin 0.5 µM, Glucose 10 mM, Oligomycin 1 µM, 2-DG 50 mM. Reagents: Seahorse XF Cell Mito Stress Test Kit 102720-100, Seahorse XF DMEM assay medium pack pH 7.4 103680-100, Seahorse XFe96 FluxPak mini 102601-100. After assay, cells were lysed in RIPA buffer followed by Bradford assay to determine total protein levels for normalization. Analysis was performed with Agilent Seahorse Wave software.

### Mitochondria Isolation

1.5 x 10^7^ cells were collected for mitochondria isolation using kit from ThermoFisher Scientific (89874), with small adjustments to centrifugation times to quickly isolate mitochondria for experimentation. Halt protease Inhibitor cocktail was added to all solutions (Thermo Fisher Scientific 87786). After spinning down for 5 min at 1000 RPM, cells were resuspended in Reagent A and split into two eppendorf tubes. Cells were lysed with pestle (VWR 47747-358) with 50 strokes on ice. After lysis, cells were centrifuged at 700 x g for 5 min, followed by mitochondria collection. Samples were combined, followed by second centrifugation for mitochondria pellet at 3000 x g for 10 min. Mitochondria pellet was washed and spun at 12,000 x g for 5 min. Mitochondria were resuspended in either PBS or Neuronal Media, followed by Bradford assay to assess total protein. For 96-well plate, mitochondria were seeded at 5 µg per well. For MEA, mitochondria were seeded at 20 µg per well.

### Western Blots

Cells were lysed in RIPA buffer (Thermo Fisher Scientific 89900) with Halt Protease Inhibitor Cocktail, and subject to standard immunoblotting analysis. Primary antibodies MECP2 anti-rabbit Invitrogen 49-1029, Beta-Actin anti-mouse, GAPDH abcam ab8245, VDAC anti-rabbit D73D12 Cell Signal Technology, TOMM20 anti-rabbit. Secondary antibodies IRDye 680RD anti-mouse Licor 926-68070, IRDye 800CW anti-rabbit Licor 926- 32211. Note - we changed the fluorescent signal to black and white in Image Studio.

### Multielectrode Array Recording

1 x 10^5^ NGN2 hESC were plated and matured at least 2 months in Matrigel plus laminin coated 48-well CytoView plates (Axion Biosystems # M768-tMEA-48B). Recordings of spontaneous activity were taken over 30-min periods on the Maestro system (Axion Biosystems). AxIS software compiled the data collected from recordings. Data were collected for LFPs (firing frequency in Hz), electrographic burst events (minimum 3 LFPs/100 ms) and relative network activity (minimum 3 LFPs detected simultaneously between a minimum of two electrodes). LFP detection was filtered at 6 × standard deviation to remove potential artifacts.

### CLEM

#### Preparation of organoids for Light Microscopy

At Day 100, organoids were labeled with live dyes. Both dyes were added simultaneously: 100 uM NeuO (STEMCELL Tech) and 100 mM Sulforhodamine 101 (Sigma) in organoid media for 1 Hour in incubator shaking.

Organoids were transferred from culture media and fixed in 4% paraformaldehyde in 0.1M sodium cacodylate buffer for 1 hour at 4°C. Organoids were rinsed in 0.1M cacodylate and embedded in warmed 2% low-melting point agar in the cacodylate buffer. Once cooled to 4°C, 150 μm sections of organoids were prepared by vibratome, and transferred to coverslips. An additional drop of warmed agar was placed on top of the tissue to immobilize sections for handling during light microscopy and processing for resin embedding. Surrounding agar was trimmed to an asymmetrical trapezoid to maintain orientation throughout processing and imaging.

#### Light Microscopy

Light microscopy was performed on a Zeiss LSM 880 Airyscan upright confocal microscope. Sections remained adhered to coverslips during imaging, and were kept wet by adding DI water when needed. Regions of organoid slices were identified with a mix of cell populations and organized morphology using a 40x water- immersion 1.1NA objective. Z-stacks were collected at 0.1 μm x-y pixel size with 0.54 μm slice spacing, and a subsequent wider 1.5 μm slice spacing to include the tissue margins. Regional z-stacks were acquired by 20x 0.8NA objective at 0.15 μm x-y pixel size and 2.2 μm slices, and overview widefield images in transmitted light and fluorescent channels were acquired by 10x 0.3NA objective lens.

#### Tissue processing for Electron Microscopy

Agar-immobilized organoid sections were post-fixed in 2.5% glutaraldehyde in 0.1 M cacodylate buffer, with special care to keep the exposed side of tissue in contact with solutions. Tissue processing for resin embedding was a variation on a ROTO protocol (Tapia et al., 2012^71^). In short, samples were further fixed in 1% osmium tetroxide 1.25% potassium ferrocyanide in 0.1 M cacodylate buffer for 1 hour at 4°C, incubated in 1% aqueous thiocarbohydrazide for 20 minutes at room temperature, and finally fixed in 1% aqueous osmium tetroxide for 1 hour at 4°C. Samples were in bloc stained overnight in 2% uranyl acetate in 0.05 M Maleate buffer. Dehydration was performed in a graded series of ethanol followed by transition into propylene oxide and infiltration into EPON 812 epoxy resin. Slices were oriented in a flat-bottomed mold with tissue facing the block face and embedded in fresh EPON 812. Blocks were polymerized at 60°C for 48 Hours.

#### Ultramicrotomy

Polymerized blocks were trimmed by hand to the extent of the tissue slice, and serial semi thin sections (0.5 μm) were collected on glass slides, stained with toluidine blue, and observed by light microscopy to locate the region of interest. Locations of nuclei and tissue margins were used to estimate and adjust sectioning orientation and depth with respect to the fluorescent z-stacks. When a semi-thin section contained a portion of the region of interest, 60nm thin sections were collected on TEM grids. Thin bar hexagonal mesh copper grids coated in a carbon-stabilized nitrocellulose film were used to expose as much of the section area as possible on the grid, while also minimizing section distortions and motions during exposure to the electron beam. After grid collection for STEM imaging, an additional semithin section is collected on a glass slide, coverslipped, and a montage acquired using a 63x oil-immersion 1.4NA objective as an alignment aid during imaging.

#### STEM imaging

Electron microscopy was performed on a Zeiss Crossbeam 540 scanning electron microscope equipped with an annular STEM detector. Data was acquired at 30kV and 362pA probe current, using the ATLAS5 automation software. Tissue was located on the grids at low magnification and 100nm x-y pixel resolution overview mosaics of the whole tissue region were acquired for each sample condition. Fluorescent, transmitted light, and semithin section data were imported into the ATLAS5 software and co-aligned with the STEM overview data to aid in navigation and correct for compression during sectioning. Regions of interest were identified by intrinsic landmarks within the organoids, and final alignment of the fluorescent z-stacks in 3D was performed through identification of multiple nuclei and cell margins. Using the fluorescent overlay a STEM dataset of 4nm x-y pixel resolution mosaics comprising the original light microscopy field of view was acquired.

#### Mitochondria Quantification

30 images were selected per genotype for imaging analysis, where images were selected with clearly defined mitochondria in regions labeled for neurons or glia. Imaging analysis was performed in Image J Fiji utilizing Region of Interest tools according to Lam et al. *Cells* 2021^72^.

### Multiomics Analysis

#### Organoid Preparation and Sequencing

At Day 100, organoids were flash frozen and stored in -80 °C. We followed the 10X Genomics Chromium Next GEM Single Cell Multiome ATAC + Gene Expression (1000285) protocol. Nuclei isolation was followed according to protocol (CG000375 Rev B). Organoids were homogenized with pestle (VWR 47747-358) with 50 strokes and incubated on ice for 5 min. After checking that cells were completely lysed, nuclei were counted with EVE automated cell counter (NanoEnTek). Nuclei were not sorted to maintain integrity. Equal number of nuclei were loaded (4500) according to protocol CG000338 Rev A.

#### Data Analysis

Libraries were sequenced on a NovaSeq 6000. We used the 10X Genomics Cell Ranger ARC software v2.0.1, “cellranger-arc mkfastq” to make the fastq files, and “cellranger-arc count” to align the reads, assign them to cells, remove duplicates and count. We used the R packages Seurat, version 4.1.1.9001, and Signac, version 1.6.0, to process and integrate the multiomics data. We followed the multiomic analysis vignette from Signac (https://stuartlab.org/signac/articles/pbmc_multiomic.html). For each or the four samples we made Seurat objects containing the gene expression data. Chromatin assays, containing the ATAC-seq data, were added to the respective Seurat objects. We merged the four objects into one Seurat object. We calculated the nucleosome signal for each cell barcode and the TSS enrichment score with the “NucleosomeSignal()” and TSSEnrichment() functions respectively. We kept cell barcodes that had a nCount_RNA between 2000 and 100000, a nCount_ATAC between 400 and 600000, nucleosome signal lower than 2, and a TSS enrichment score higher than 1. After this filtering we retained 1914 cells from the control 1 organoid, 1722 from the control 2 organoid, 2901 cells from the RTT 1 organoid, and 747 cells from the RTT 2 organoid. We defined peaks with MACS2 using the ATAC-seq fragments from all the samples. We quantified the fragments overlapping with the peaks, and made a new chromatin assay, “peaks assay” that we used for the ATAC-seq analysis. We integrated the samples on the gene expression assays using cluster similarity spectrum (CSS)^73^ implemented in the R package simspec, following the vignette (https://github.com/quadbiolab/scRNAseq_analysis_vignette/blob/master/Tutorial.md#step-2-6-data-integration-using-css). We processed and integrated the peaks assay following the scATAC-seq data integration vignette from Signac, (https://stuartlab.org/signac/articles/integrate_atac.html). Briefly, we used the default Signa implementation of the term frequency-inverse document frequency (TF-IDF) normalization, feature selection and singular value decomposition (SVD) on the TF-IDF matrix. The result of these steps is known as latent semantic indexing (LSI)^74^. We run uniform manifold approximation and projection (UMAP) on the merged Seurat objects using lsi components 2 to 30 as input, since the first lsi component correlated with sequencing depth. To integrate the samples on the peak assays, we selected the integration anchors from peaks that were at 1 Kb or less from a protein coding gene with the “FindIntegrationAnchors” function setting the reduction parameter to "rlsi" and using dimensions 2 to 30. We then integrated the datasets using the “IntegrateEmbeddings“ function, the anchors selected on the previous step and the 1 to 30 lsi reductions calculated for each sample. We ran UMAP on the resulting integrated lsi using dimensions 1 to 30. Finally, we combined the integrated expression and peak assays via weighted nearest neighbor (WNN) analysis running the function “FindMultiModalNeighbors” on the ccs (from the integrated expression assay) and integrated lsi (from the integrated peaks assay) dimensionality reductions. We ran UMAP and Louvain clustering with 0.2 resolution on the weighted combination of expression and ATAC-seq data. We found 11 clusters. We identified genes specific of each cluster with the “FindAllMarkers“ function, selecting only positive markers. To assign cell identities to the clusters, we used the cluster specific markers and the cell-type specific expression analysis tools developed by Camp et al.^35^ and Fleck et al.^36^, a human cortex dataset from Nowakowski et al.^37^, and the BrainSpan Atlas of the Developing Human Brain^38^ via Cell-type Specific Expression Analysis tool developed by Dougherty *et al*.^39^.

We converted the Seurat object into a single cell object and exported the sum of the raw gene or peaks counts for the groups of cells we wanted to compare, *e.g*., “CRT1” cells or “CTR1 neuronal” cells. We proceeded to analyze these pseudo-bulk counts matrices with DESeq2^75^. In the absence of spike ins, the recommended method for normalizing ATAC-seq data is the use of “full library size factors”^76^. These are normalization factors derived from the total number of fragments in each library, include background reads, and would not be affected by changes in a subset of the opened regions. We normalized our ATAC-seq peaks count matrix using the DESeq2 default, total number of fragments in peaks, as well as using full library size factors. Both sets of factors were very close: CTR1: 0.89, CTR2: 1.29, RTT1: 1.67, and RTT2: 0.50 using counts in peaks, and CTR1: 0.83, CTR2: 1.19, RTT1: 1.53, and RTT2: 0.45, using full library counts. The output from both methods was equivalent. This is not unexpected since we did not filter out any low confidence peaks that would include mostly background reads. To normalize the pseudo bulk RNA-seq from the single nuclei expression we used the standard DESeq2 normalization method. This method calculates a size factor from the total number of reads mapped to genes, and removes any global changes in gene expression present in the data. Since global gene repression has previously been reported for mutant MECP2, we aimed to detect such global change by normalizing the RNA-seq data with size factors independent of the gene expression count matrix. We used the full library size factors from the ATAC-seq library and found a small global gene repression as previously reported^19, 65^. To further confirm that the increase in expression of specific genes was not affected by this global repression, we lowered the log2 fold change (log2FC) for each gene by the median log2FC previously reported^65^, log2FC-0.1874, and found this shift to have a negligent impact in our conclusions. The figures and subsequent analysis for both pseudo-bulk ATAC-seq and RNA-seq were done using the DESeq2 default calculation of size factors derived from the feature count matrix. We did not shrink the log2FC of the ATAC-seq peaks analysis. We used normal shrinkage when analyzing the expression data. We ran gene set enrichment analysis with GSEA 4.3.2^77^ ranking the genes by the statistic from the DESeq2 Wald test. We used Cluster 3.0^78^ the R library heatmap and a custom script to make the heatmap figures.

### Statistical Analysis

Statistical parameters including sample sizes and statistical significance are reported in the Figures and the Figure Legends. Data is judged to be statistically significant when p ≤ 0.05 by two-tailed Student’s T-Test, One-way or Two-way ANOVA, where appropriate. Statistical analysis was performed in GraphPad Prism, except for multiomics analysis described above.

## Reporting summary

### Data availability

Further information and requests for reagents may be directed to, and will be fulfilled by the corresponding author, Dr. Rudolf Jaenisch.

## Contributions

DLT and RJ designed the study and wrote the manuscript. MIB performed multiomics analysis. DM and AL-J devised and performed CLEM imaging. KIA assisted in qPCR studies, sholl and mitochondrial analysis. All other experiments and analysis were performed by DLT.

## Acknowledgements

We thank all members of the Jaenisch lab for helpful discussions and manuscript editing, reagent maintenance and management, and purchasing and administrative assistance. We thank Anthony Flamier for plasmids used for cortical neuron differentiation, Tenzin Kunchok and the Metabolite Profiling Core Facility at the Whitehead Institute for running metabolomics samples and data analysis, Fabian Schulte, Brooke Linnehan and the Quantitative Proteomics Core at the Whitehead Institute for running proteomics samples and data analysis, Jennifer Love, Sumeet Gupta, and the Genome Core at the Whitehead Institute for running multiomics samples and data analysis, Cassandra Rogers and Brandyn Braswell and the Keck Imaging Facility at the Whitehead Institute for help with high throughput imaging analysis, and Yuchen Wang for aiding in mitochondrial analysis. We thank Mike Gallagher and Max Friesen for helpful comments while assembling the manuscript. This work was supported in part by the Koch Institute Support (core) Grant P30-CA14051 from the National Cancer Institute. We thank the Koch Institute’s Robert A. Swanson (1969) Biotechnology Center for technical support, specifically Peterson (1957) Nanotechnology Materials Core Facility (RRID:SCR_018674)

## Extended Data Figures

**Extended Data Fig. 1.**
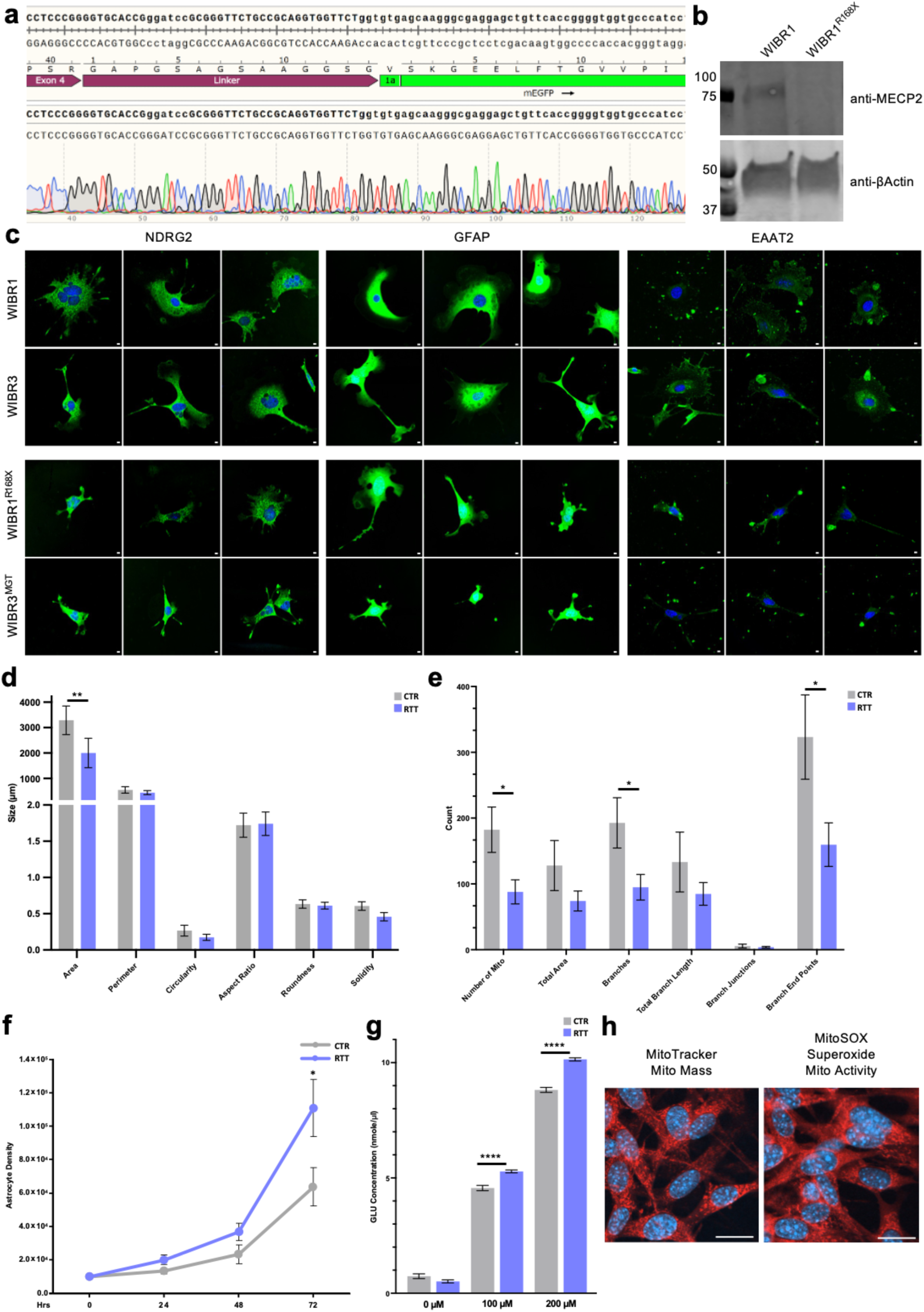
**a)** Plasmid pCL186_pUC19_HDR_. CRISPR. Sanger sequencing result confirmed that a linker-eGFP transgene is inserted after MECP2-R168 in the Exon 4 position to result in early truncation. **b)** Western blot confirming absence of MECP2 C-terminus in mutated WIBR1^R168X^ hESC compared to WIBR1 CTR. Truncated MECP2 fused with GFP predicted similar size as full length MECP2 **c)** Representative images of ASTs seeded at a low density derived from CTR hESC WIBR1 and WIBR3 (top panel) and RTT hESC WIBR1^R168X^ and WIBR3^MGT^ (bottom panel). Expression of AST markers NDRG2, GFAP, and EAAT2 were used to define maturity. Merge with DAPI. Scale bar 5 µm. **d)** Quantification of AST parameters with Beta-Actin staining from Fig. 1a. Statistical analysis Two-Way ANOVA Bonferroni **p ≤ 0.01 Error bars SEM. **e)** Quantification of mitochondria parameters with TOM20 staining from Fig. 1a. Statistical analysis individual T-Test *p ≤ 0.05. Error bars SEM. **f)** AST proliferation over 72 hours (hrs). 24 hrs CTR n = 12, RTT n = 12; 48 hrs CTR n = 16, RTT n = 16; 72 hrs CTR n = 19, RTT n = 20. Statistical analysis One-Way ANOVA Tukey *p ≤ 0.05. Error bars Standard Error of Measure (SEM). **g)** Glutamate uptake assay. Extracellular glutamate was measured after 4 hr incubation. 0 µM CTR n = 16, RTT = 24, 100 µM CTR n = 16, RTT n = 24, 200 µM CTR n = 30, RTT n = 54. Statistical analysis Two-Way ANOVA Multiple Comparisons Bonferroni ****p ≤ 0.0001. **h)** Representative images of MitoTracker (top) or MitoSOX (bottom) staining. Scale bar 10 µM.

**Extended Data Fig. 2.**
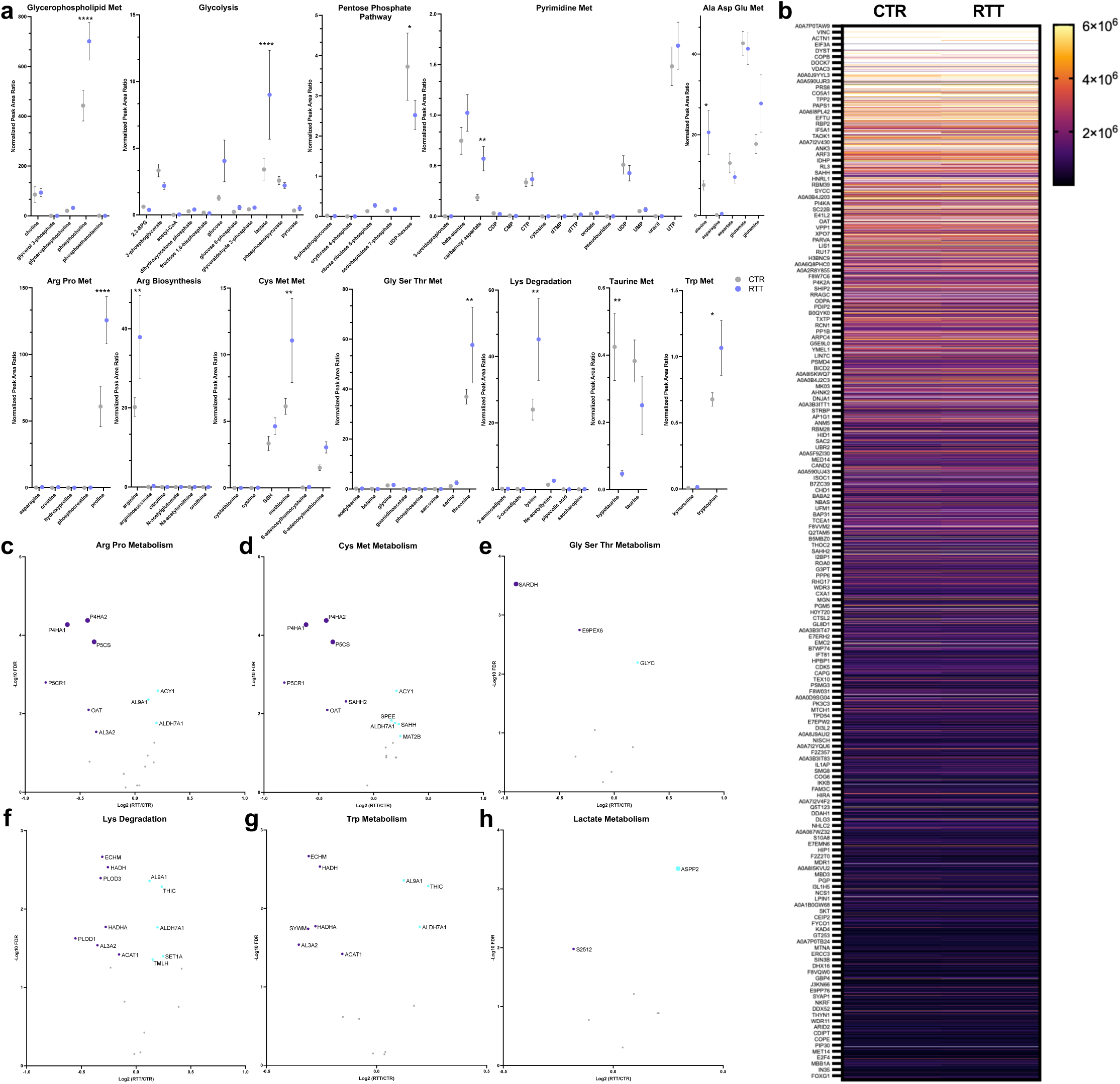
**a)** Whole-cell AST polar metabolite analysis, only statistically significant pathways indicated. CTR n = 9, RTT n = 10. Statistical analysis Two-Way ANOVA Multiple Comparisons Bonferroni *p ≤ 0.05, **p ≤ 0.01, ***p ≤ 0.001, ****p ≤ 0.0001. Error bars SEM. Non-significant pathways not shown (Cofactor/Redox Metabolism, Fatty Acid Synthesis, One-Carbon Metabolism, Purine Metabolism, Histidine Metabolism, Isoleucine Leucine and Valine Metabolism, Phenylalanine Metabolism). **b)** Heatmap of all proteins from proteomic analysis from Fig. 3. Every 20th protein labeled. **c-h)** Volcano plots of whole cell AST proteomic analysis (Fig. 3). Protein changes in specific metabolic pathways for arginine and proline (**c**), cysteine and methionine (**d**), glycine, serine and threonine (**e**), lysine degradation (**f**), tryptophan (**g**) and lactate (**h**). Increased proteins (light blue), decreased proteins (purple) in RTT compared to CTR. Large dot adjusted ***p ≤ 0.001, small dot adjusted *p ≤ 0.05. CTR n = 6, RTT n = 6.

**Extended Data Fig. 3.**
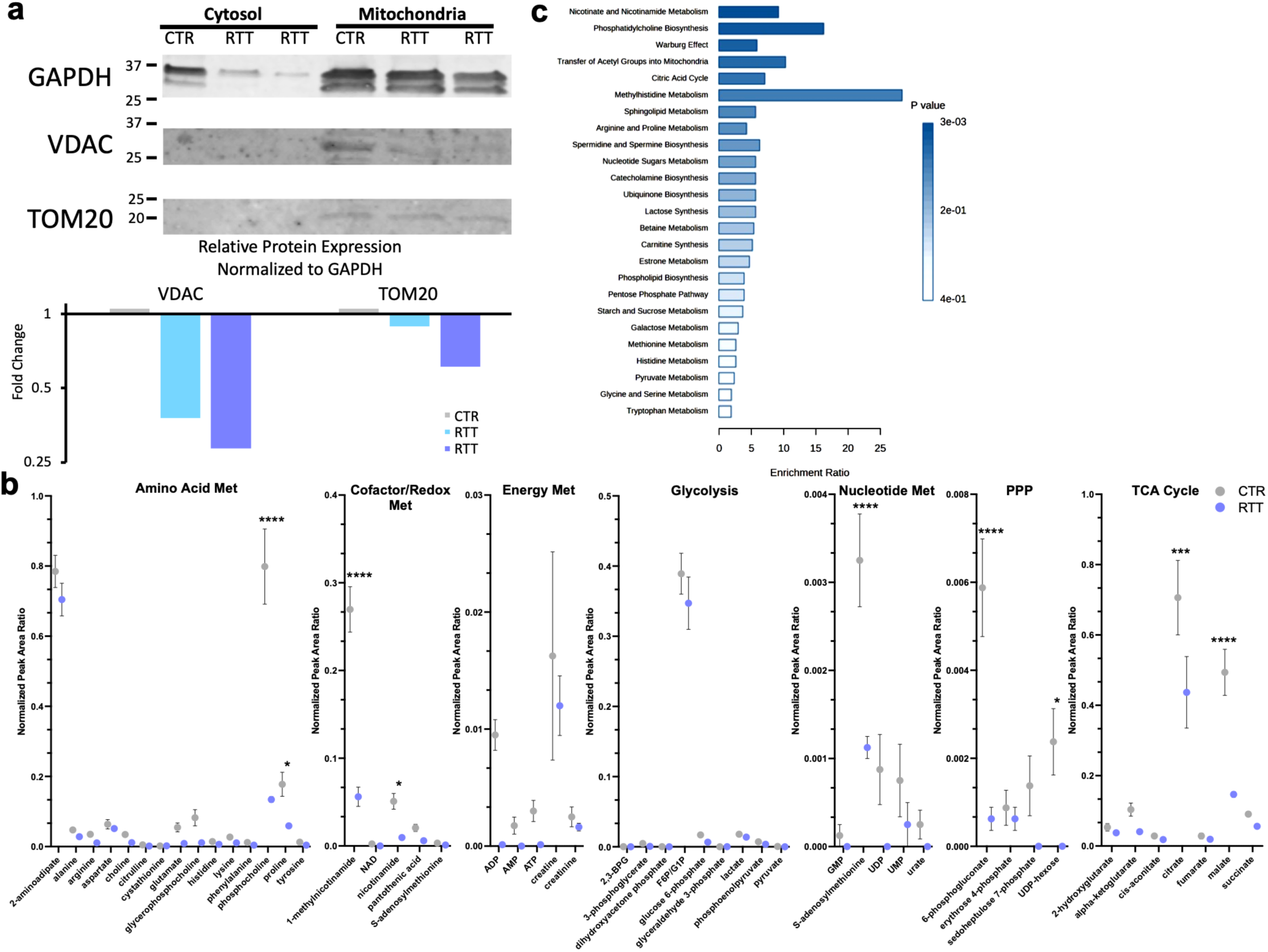
**a)** Western blot of cytosolic and mitochondria fractions after mitochondria isolation. Presence of mitochondrial membrane proteins VDAC and TOM20 indicates separation of organelle. Fold change of WIBR1^R168X^ and WIBR3^MGT^ RTT relative to WIBR1 CTR after densiometric analysis normalized to GAPDH. **b)** Isolated mitochondria from AST polar metabolite analysis. CTR n = 6, RTT n = 6. Statistical analysis Two-Way ANOVA Multiple Comparisons Bonferroni *p ≤ 0.05, ***p ≤ 0.001, ****p ≤ 0.0001. Error bars SEM. **c)** Metabolism enrichment analysis of mitochondrial metabolites (**b**).

**Extended Data Fig. 4.**
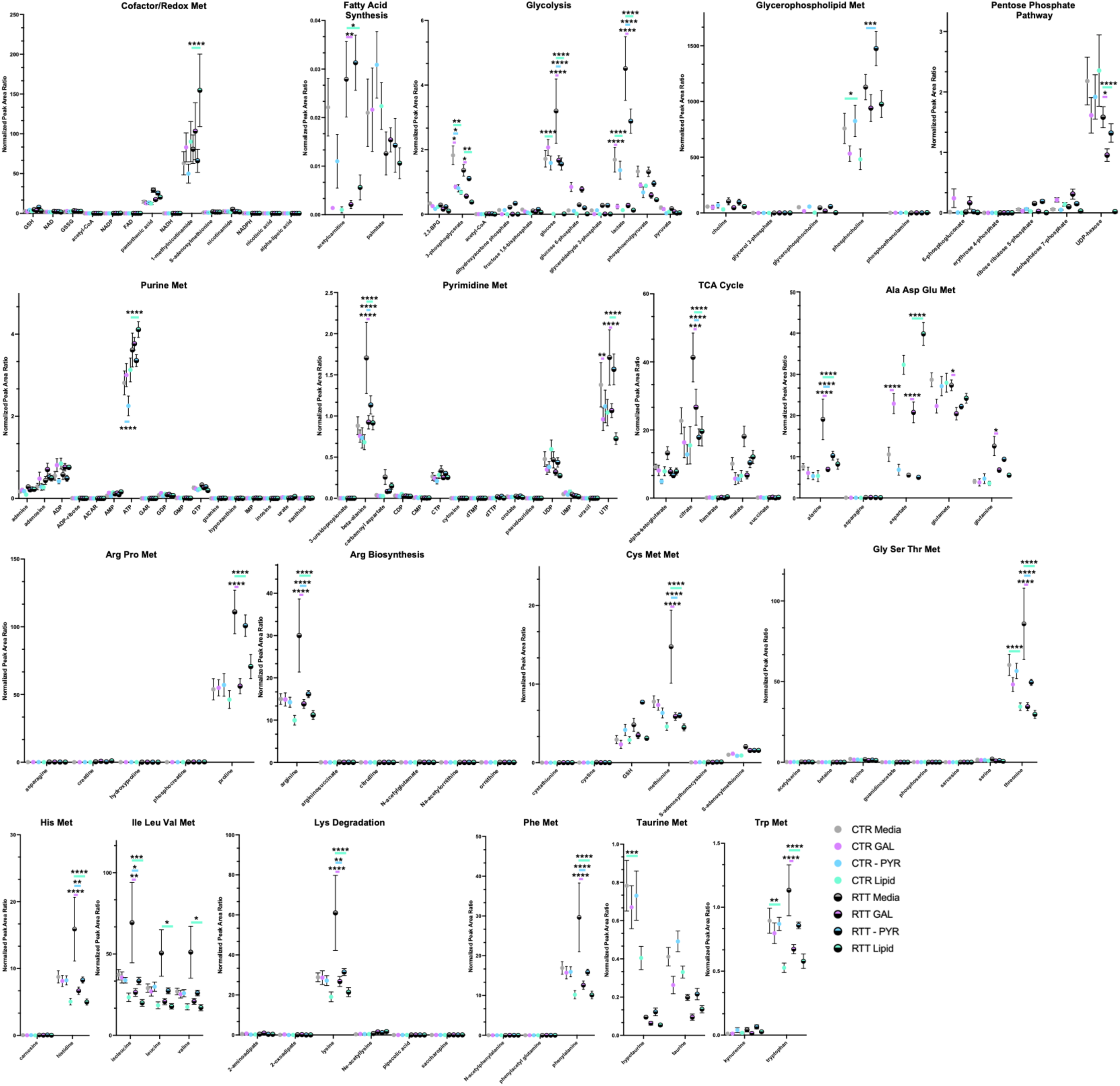
Whole cell AST metabolomic analysis. Incubation with conditions for 24 hrs. Media (Glucose), GAL (Galactose), -PYR (Glucose minus sodium pyruvate), LIP (Lipid cocktail). Biological replicates CTR n = 8, CTR GAL n = 8, CTR –PYR n = 8, CTR LIP n = 8, RTT n = 10, RTT GAL n = 10, RTT –PYR n = 10, RTT LIP n = 10. Statistical analysis Two-Way ANOVA Multiple Comparisons Bonferroni *p ≤ 0.05, **p ≤ 0.01, ***p ≤ 0.001, ****p ≤ 0.0001. Error bars SEM.

**Extended Data Fig. 5.**
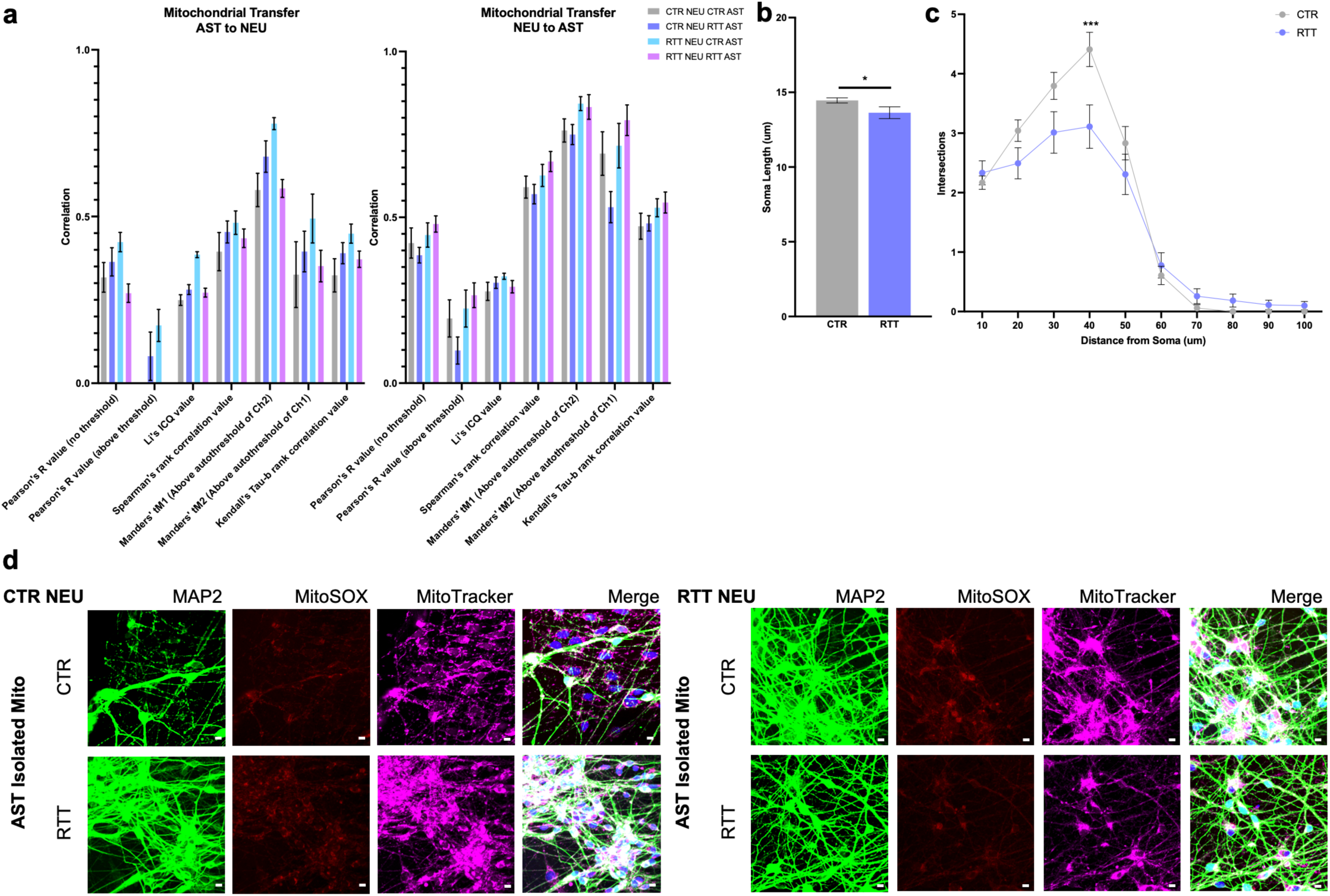
**a)** Colocalization analysis from Fig. 5a. Channel 1 – MAP2 or GFAP, Channel 2 – MitoTracker. Statistical analysis 2-Way ANOVA comparing transfer of mitochondria from AST to NEU to NEU to AST. Pearson’s R value (No threshold): RTT NEU RTT AST *p ≤ 0.05. Pearson’s R value (Above threshold): CTR NEU CTR AST **p ≤ 0.01, RTT NEU RTT AST ****p ≤ 0.0001. Spearman’s rank correlation value: RTT NEU RTT AST *p ≤ 0.05. Mander’s tM1: RTT NEU RTT AST **p ≤ 0.01. Mander’s tM2: CTR NEU CTR AST ****p ≤ 0.0001, RTT NEU RTT AST ****p ≤ 0.0001. Error bars SEM. **b)** Soma Size. Statistical analysis Unpaired T-Test. CTR n = 192, RTT n = 81. Error bars SEM. **c)** Sholl analysis. Statistical analysis Two-Way ANOVA Multiple Comparisons Bonferroni ***p ≤ 0.001. Error bars SEM. **d)** Mitochondria transplantation assay. Isolated mitochondria from RTT and CTR ASTs (left column) were labeled with Mitotracker and seeded onto CTR and RTT cortical neurons. Cultures were incubated for 48 hrs before fixing and staining. CTR and RTT cortical neurons can take up both CTR and RTT mitochondria. Scale bar 5 µm.

**Extended Data Fig. 6.**
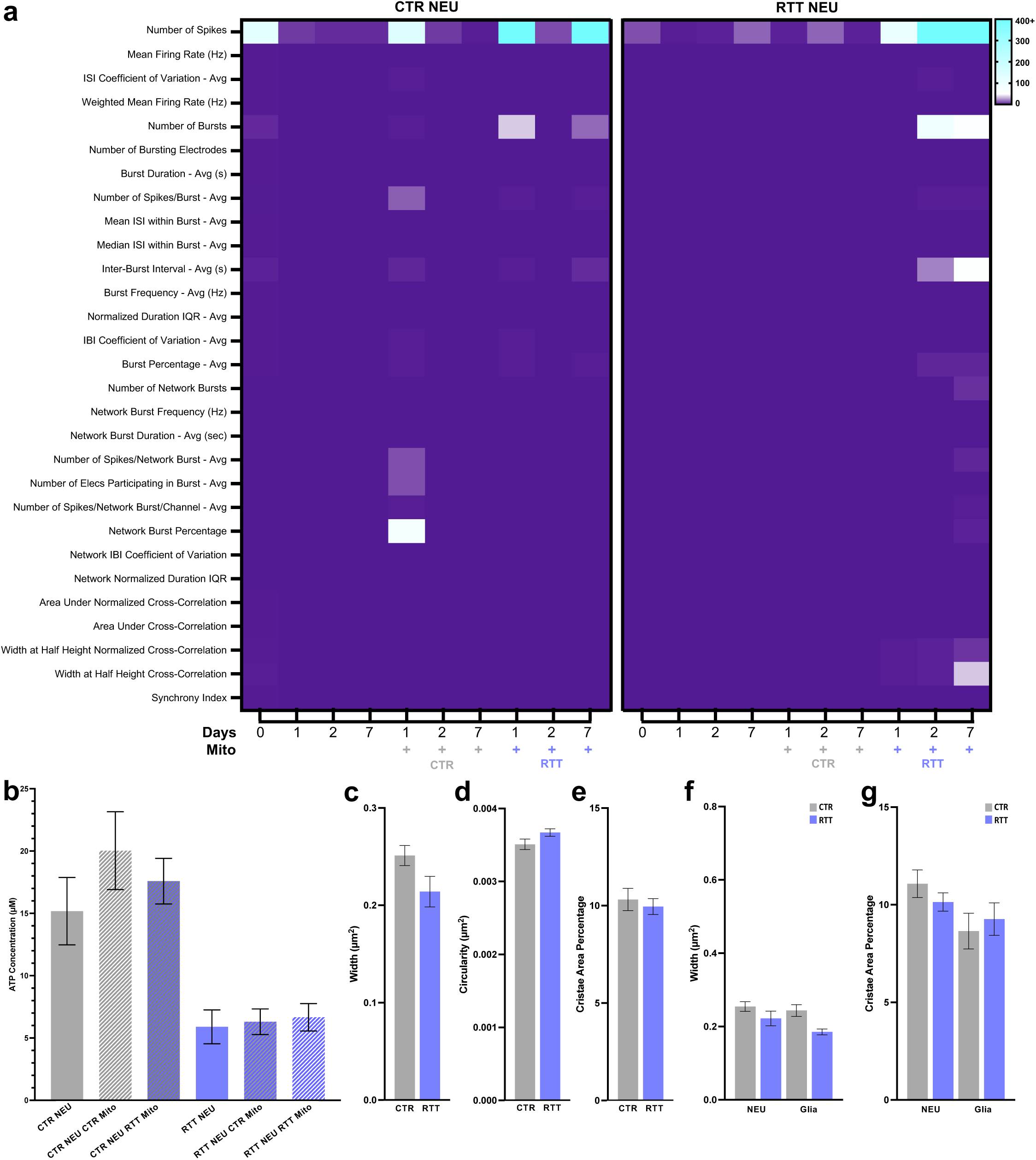
**a)** Heatmap of MEA parameters from Fig. 5c**. b)** Total neuronal ATP levels after 48 hr incubation with or without exogenous mitochondria isolated from CTR and RTT ASTs. No significant difference within CTR and RTT groups after mitochondria were seeded for 48 hrs by Two-Way ANOVA Bonferroni. Error bars SEM. **c-g)** Quantification of mitochondria ultrastructure morphology. Total CTR n = 251, total RTT n = 430, NEU CTR n = 172, NEU RTT n = 337, Glia CTR n = 79, Glia RTT n = 93.

**Extended Data Fig. 7.**
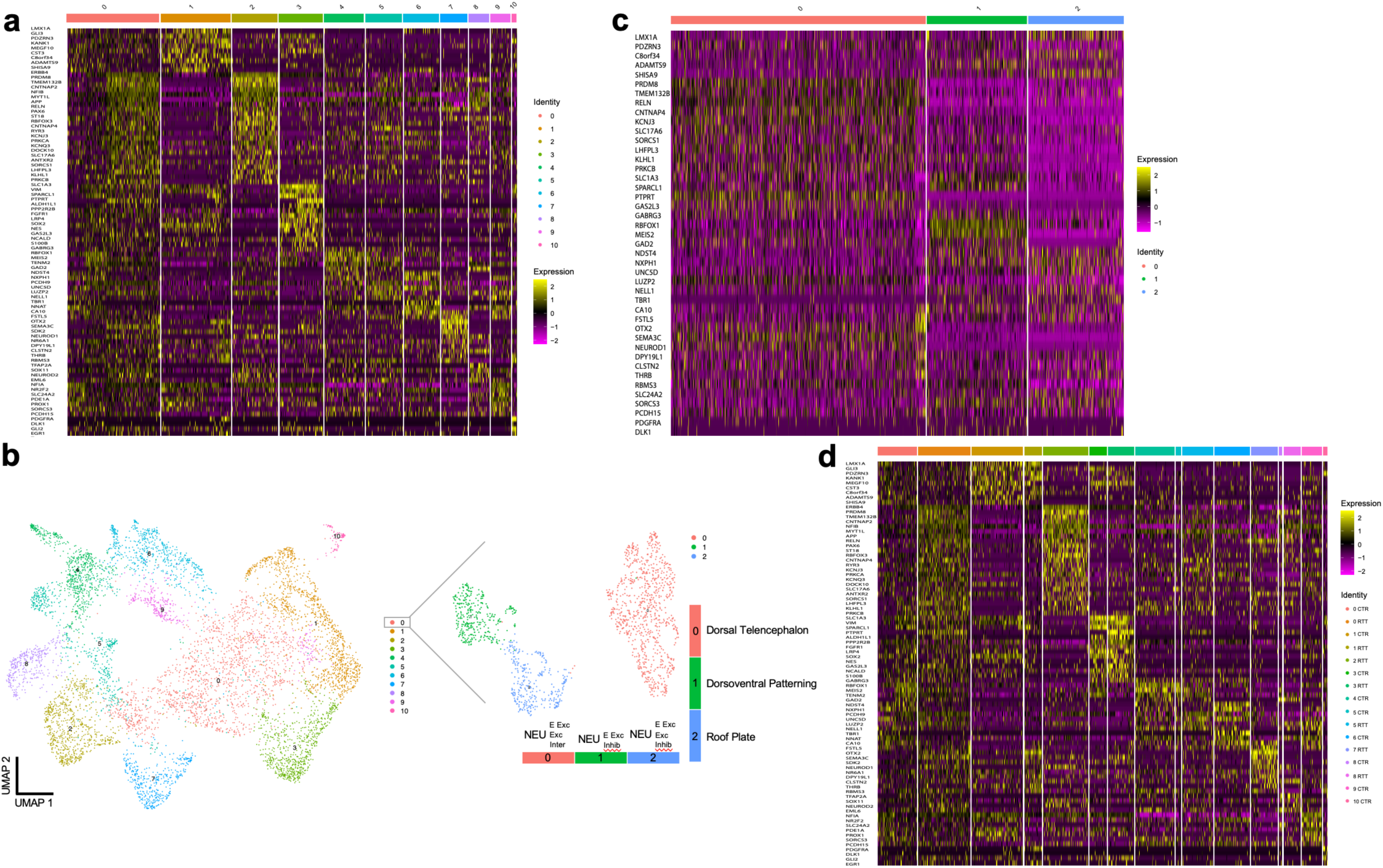
**a)** Heatmap of top genes that defined clusters in Fig. 6b. **b)** Subclustering of cluster 0 from Fig. 6b. Bottom legend cell identities: Neuron (NEU), Early Excitatory NEU (E Exc), Excitatory NEU (Exc), Interneuron (Inter), Inhibitory NEU (Inhib). Right legend early brain anatomy. **c)** Heatmap of top genes that defined clusters in (**a**) for cluster 0 subclusters. **d)** Heatmap of clusters separated by genotype.

**Extended Data Fig. 8.**
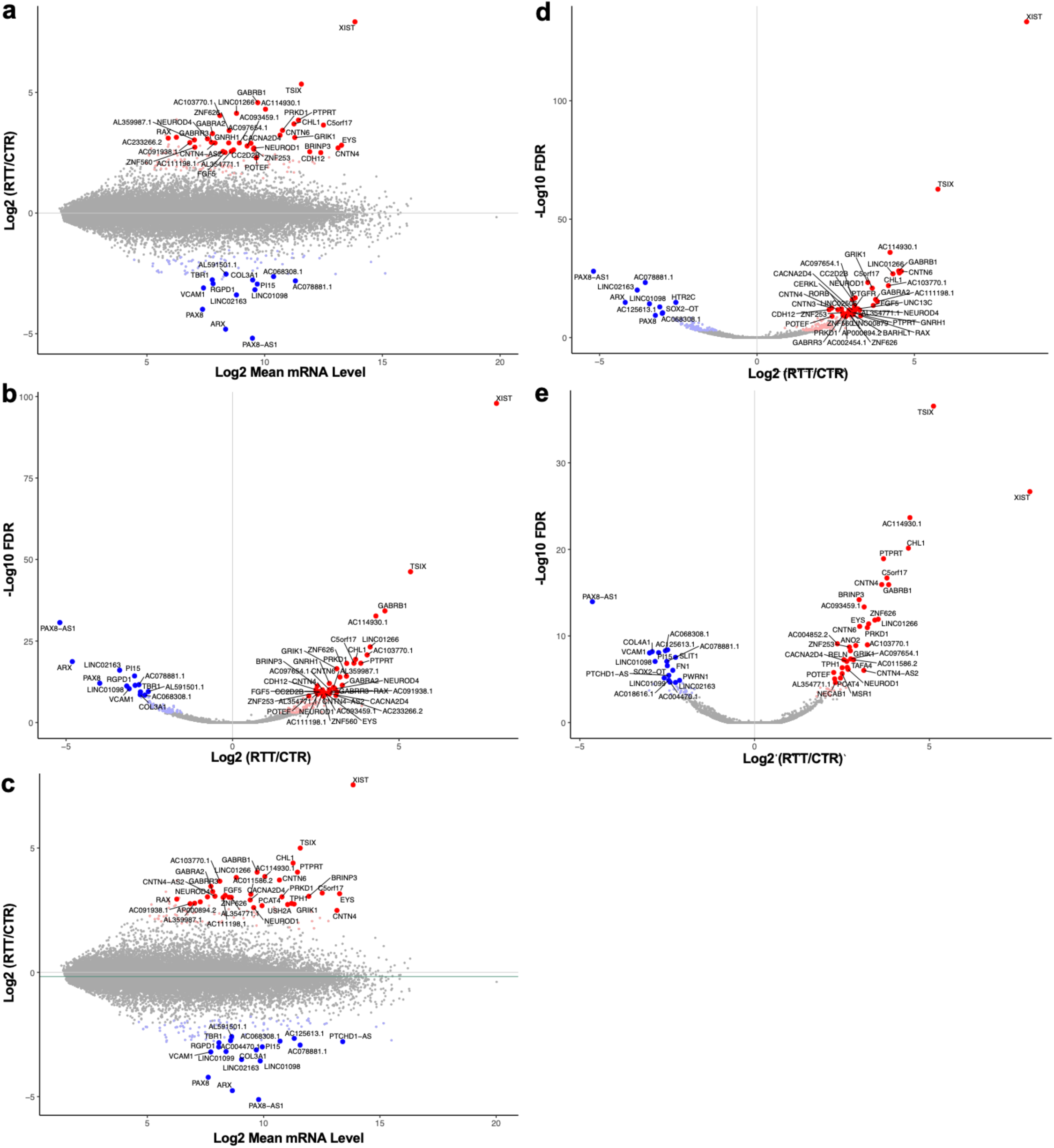
**a)** MA plot total gene expression changes from snRNA-seq of RTT compared to CTR using DESeq2 default normalization. **b)** Volcano plot of differential gene expression changes from snRNA-seq of RTT compared to CTR. **c)** MA plot total gene expression changes from snRNA-seq of RTT compared to CTR using the size factors derived from the full library size factors from the ATAC-seq library. Green line indicates shift from zero of the median fold change. **d)** Volcano plot neuronal clusters, RTT compared to CTR. **e)** Volcano plot of glial clusters, RTT compared to CTR. Significantly increased transcripts in red, significantly decreased transcripts in blue. Adjusted ***p ≤ 0.001. Top 50 genes are labelled.

**Extended Data Fig. 9.**
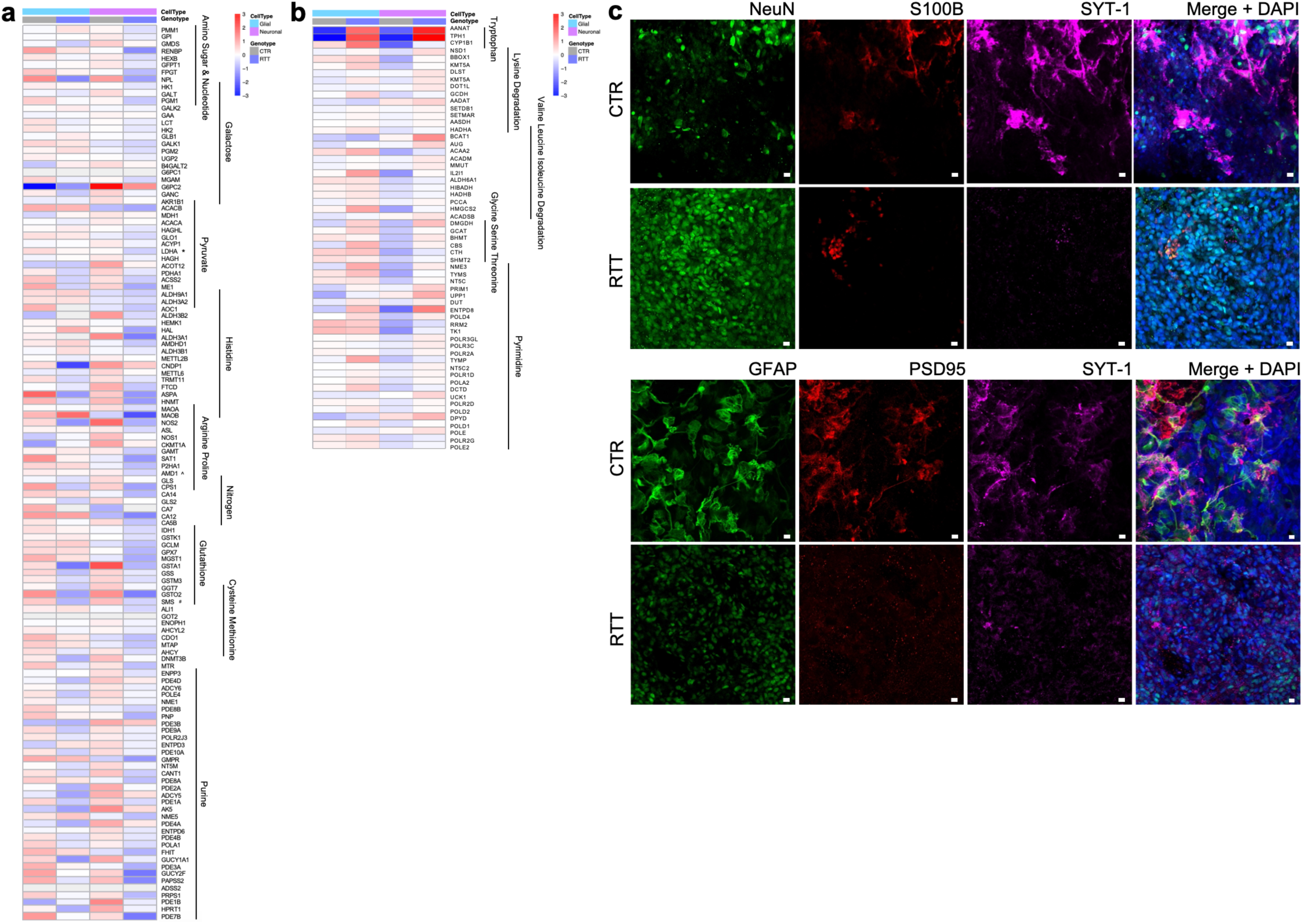
**a)** Heatmap of combined KEGG Metabolism GSEA sets enriched in neuron CTR population compared to neuron RTT population. Genes within sets (identified on right) were combined. **b)** Heatmap of combined KEGG Metabolism GSEA sets enriched in neuron RTT population compared to neuron CTR population. Genes within sets (identified on right) were combined. *LDHA – also included in Cysteine Methionine Metabolism. ^AMD1 – also included in Cysteine Methionine Metabolism. #SMS - also included in Arginine Proline Metabolism. **c)** Representative images of Day 100 cerebral organoids immunohistochemistry. NeuN (neuronal marker), SYT-1 (presynaptic marker), PSD-95 (post-synaptic marker), S100B and GFAP (AST markers). Merge with DAPI, scale bar 5 µm.

**Extended Data Fig. 10.**
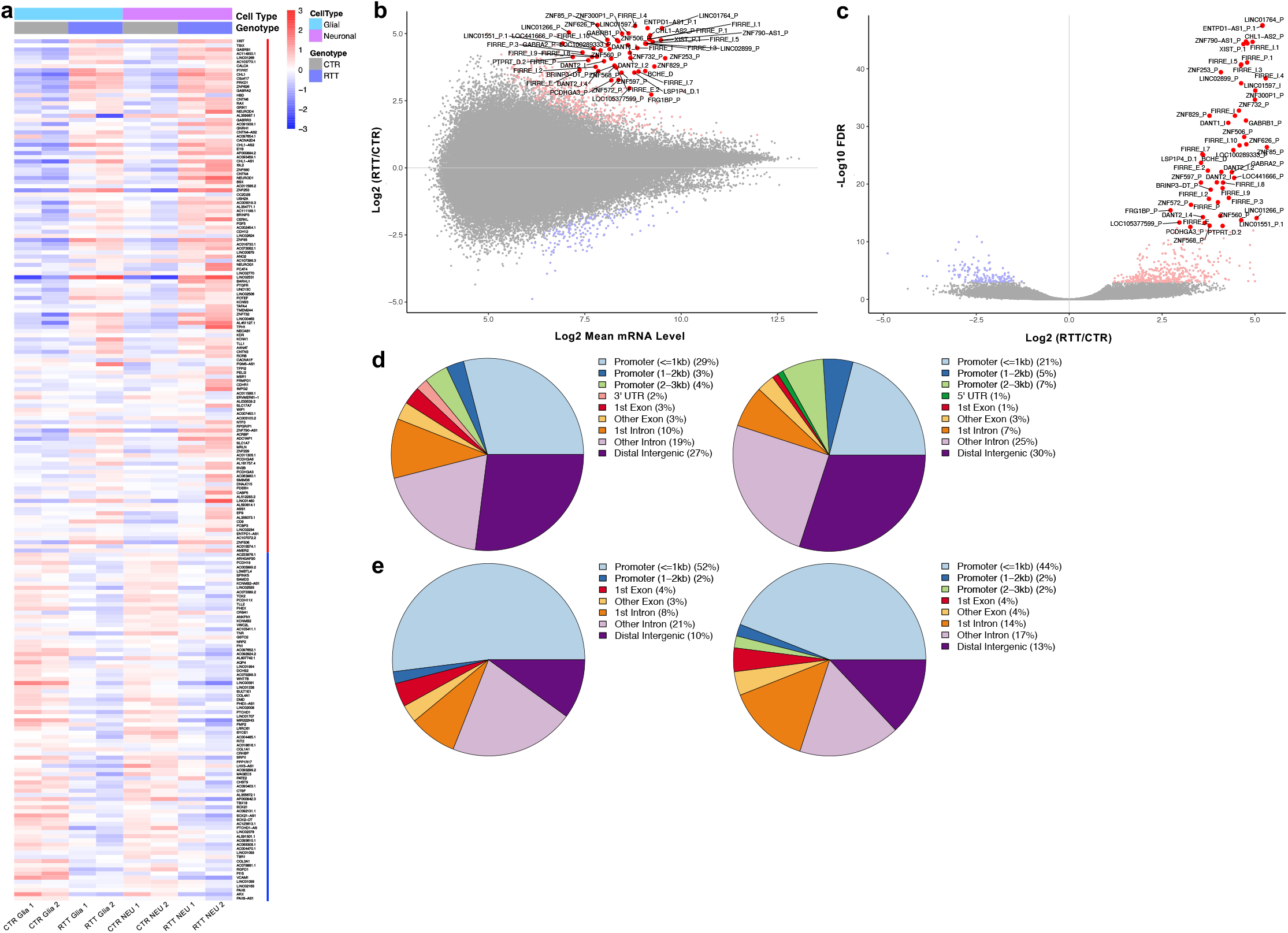
**a)** ATAC-seq normalized signal within promoter regions (3Kb upstream and 1kb downstream of the gene start) of top differentially expressed genes in RTT compared to CTR. Red line - increased in RTT, blue line - decreased in RTT compared to CTR. Adjusted ***p ≤ 0.001. **b)** MA plot total ATAC peak accessibility changes from snATAC-seq of RTT compared to CTR. **c)** Volcano plot of differentially accessible regions from snATAC- seq in RTT compared to CTR. **b-c)** Letter after gene describes position of ATAC peak – Promoter (P), Intron (I), Distal Intergenic (D), Exon (E). Significantly increased transcripts in red, significantly decreased transcripts in blue. Adjusted ***p ≤ 0.001. Top 50 regions are labelled. **d)** Distribution of the top 100 differentially accessible regions more accessible in CTR compared to RTT in glia (left) and neurons (right). **e)** Distribution of the top 100 differentially accessible regions more accessible in RTT compared to CTR in glia (left) and neurons (right).

**Supplementary Table 1.** Cell count distribution in clusters and cell type distribution compared to total cells. Note - genotype with less than 10 cells per cluster were removed from cluster classification.

| Cell Count | Cluster 0 | Cluster 1 | Cluster 2 | Cluster 3 | Cluster 4 | Cluster 5 | Cluster 6 | Cluster 7 | Cluster 8 | Cluster 9 | Cluster 10 | Total |
| --- | --- | --- | --- | --- | --- | --- | --- | --- | --- | --- | --- | --- |
| CTR | 655 | 863 | 7 | 291 | 654 | 96 | 598 | 0 | 59 | 333 | 80 | 3636 |
| RTT | 883 | 298 | 759 | 442 | 12 | 517 | 2 | 445 | 289 | 0 | 1 | 3648 |
| Composition | Neurons | Neuron % | Glia | Glia % |  |  |  |  |  |  |  |  |
| CTR | 2402 | 66.06 | 1234 | 33.94 |  |  |  |  |  |  |  |  |
| RTT | 2907 | 79.69 | 741 | 20.31 |  |  |  |  |  |  |  |  |

## References

1. Ip, J. P. K., Mellios, N. & Sur, M. Rett syndrome: insights into genetic, molecular and circuit mechanisms. Nature Reviews Neuroscience 19, 368–382, doi:10.1038/s41583-018-0006-3 (2018).

2. Jin, X. R., Chen, X. S. & Xiao, L. MeCP2 Deficiency in Neuroglia: New Progress in the Pathogenesis of Rett Syndrome. Front Mol Neurosci 10, 316, doi:10.3389/fnmol.2017.00316 (2017).

3. Sofroniew, M. V. & Vinters, H. V. Astrocytes: biology and pathology. Acta Neuropathol 119, 7–35, doi:10.1007/s00401-009-0619-8 (2010).

4. Okabe, Y. et al. Alterations of gene expression and glutamate clearance in astrocytes derived from an MeCP2-null mouse model of Rett syndrome. PLoS One 7, e35354, doi:10.1371/journal.pone.0035354 (2012).

5. Dong, Q., Kim, J., Nguyen, L., Bu, Q. & Chang, Q. An Astrocytic Influence on Impaired Tonic Inhibition in Hippocampal CA1 Pyramidal Neurons in a Mouse Model of Rett Syndrome. J Neurosci 40, 6250–6261, doi:10.1523/JNEUROSCI.3042-19.2020 (2020).

6. Williams, E. C. et al. Mutant astrocytes differentiated from Rett syndrome patients-specific iPSCs have adverse effects on wild-type neurons. Hum Mol Genet 23, 2968–2980, doi:10.1093/hmg/ddu008 (2014).

7. Caldwell, A. L. M. et al. Aberrant astrocyte protein secretion contributes to altered neuronal development in multiple models of neurodevelopmental disorders. Nat Neurosci 25, 1163–1178, doi:10.1038/s41593-022-01150-1 (2022).

8. Ballas, N., Lioy, D. T., Grunseich, C. & Mandel, G. Non-cell autonomous influence of MeCP2-deficient glia on neuronal dendritic morphology. Nature Neuroscience 12, 311–317, doi:10.1038/nn.2275 (2009).

9. Lioy, D. T. et al. A role for glia in the progression of Rett’s syndrome. Nature 475, 497–500, doi:10.1038/nature10214 (2011).

10. Sun, J. et al. Mutations in the transcriptional regulator MeCP2 severely impact key cellular and molecular signatures of human astrocytes during maturation. Cell Rep 42, 111942, doi:10.1016/j.celrep.2022.111942 (2023).

11. An, C. et al. Overcoming Autocrine FGF Signaling-Induced Heterogeneity in Naive Human ESCs Enables Modeling of Random X Chromosome Inactivation. Cell Stem Cell 27, 482–497 e484, doi:10.1016/j.stem.2020.06.002 (2020).

12. Krishnaraj, R., Ho, G. & Christodoulou, J. RettBASE: Rett syndrome database update. Hum Mutat 38, 922–931, doi:10.1002/humu.23263 (2017).

13. Lengner, C. J. et al. Derivation of pre-X inactivation human embryonic stem cells under physiological oxygen concentrations. Cell 141, 872–883, doi:10.1016/j.cell.2010.04.010 (2010).

14. Escartin, C. et al. Reactive astrocyte nomenclature, definitions, and future directions. Nat Neurosci 24, 312–325, doi:10.1038/s41593-020-00783-4 (2021).

15. Schousboe, A., Scafidi, S., Bak, L. K., Waagepetersen, H. S. & McKenna, M. C. Glutamate metabolism in the brain focusing on astrocytes. Adv Neurobiol 11, 13–30, doi:10.1007/978-3-319-08894-5_2 (2014).

16. Zhang, H., Menzies, K. J. & Auwerx, J. The role of mitochondria in stem cell fate and aging. Development 145, doi:10.1242/dev.143420 (2018).

17. de Moura, M. B., dos Santos, L. S. & Van Houten, B. Mitochondrial dysfunction in neurodegenerative diseases and cancer. Environ Mol Mutagen 51, 391–405, doi:10.1002/em.20575 (2010).

18. Shulyakova, N., Andreazza, A. C., Mills, L. R. & Eubanks, J. H. Mitochondrial Dysfunction in the Pathogenesis of Rett Syndrome: Implications for Mitochondria-Targeted Therapies. Front Cell Neurosci 11, 58, doi:10.3389/fncel.2017.00058 (2017).

19. Li, Y. et al. Global transcriptional and translational repression in human-embryonic-stem-cell-derived Rett syndrome neurons. Cell Stem Cell 13, 446–458, doi:10.1016/j.stem.2013.09.001 (2013).

20. Esparham, A. E. et al. Nutritional and Metabolic Biomarkers in Autism Spectrum Disorders: An Exploratory Study. Integr Med (Encinitas) 14, 40–53 (2015).

21. Frye, R. E. et al. Emerging biomarkers in autism spectrum disorder: a systematic review. Ann Transl Med 7, 792, doi:10.21037/atm.2019.11.53 (2019).

22. Chen, W. W., Freinkman, E., Wang, T., Birsoy, K. & Sabatini, D. M. Absolute Quantification of Matrix Metabolites Reveals the Dynamics of Mitochondrial Metabolism. Cell 166, 1324–1337 e1311, doi:10.1016/j.cell.2016.07.040 (2016).

23. You, X. et al. Loss of mitochondrial aconitase promotes colorectal cancer progression via SCD1- mediated lipid remodeling. Mol Metab 48, 101203, doi:10.1016/j.molmet.2021.101203 (2021).

24. Ward, N. P., Kang, Y. P., Falzone, A., Boyle, T. A. & DeNicola, G. M. Nicotinamide nucleotide transhydrogenase regulates mitochondrial metabolism in NSCLC through maintenance of Fe-S protein function. J Exp Med 217, doi:10.1084/jem.20191689 (2020).

25. Hayakawa, K. et al. Transfer of mitochondria from astrocytes to neurons after stroke. Nature 535, 551–555, doi:10.1038/nature18928 (2016).

26. Gordon, A. et al. Long-term maturation of human cortical organoids matches key early postnatal transitions. Nat Neurosci 24, 331–342, doi:10.1038/s41593-021-00802-y (2021).

27. Lewis, J. D. et al. Purification, sequence, and cellular localization of a novel chromosomal protein that binds to methylated DNA. Cell 69, 905–914, doi:10.1016/0092-8674(92)90610-o (1992).

28. Good, K. V., Vincent, J. B. & Ausio, J. MeCP2: The Genetic Driver of Rett Syndrome Epigenetics. Front Genet 12, 620859, doi:10.3389/fgene.2021.620859 (2021).

29. Gulmez Karaca, K., Brito, D. V. C. & Oliveira, A. M. M. MeCP2: A Critical Regulator of Chromatin in Neurodevelopment and Adult Brain Function. Int J Mol Sci 20, doi:10.3390/ijms20184577 (2019).

30. Tillotson, R. & Bird, A. The Molecular Basis of MeCP2 Function in the Brain. J Mol Biol, doi:10.1016/j.jmb.2019.10.004 (2019).

31. Della Ragione, F., Vacca, M., Fioriniello, S., Pepe, G. & D’Esposito, M. MECP2, a multi-talented modulator of chromatin architecture. Brief Funct Genomics 15, 420–431, doi:10.1093/bfgp/elw023 (2016).

32. Lee, W., Kim, J., Yun, J. M., Ohn, T. & Gong, Q. MeCP2 regulates gene expression through recognition of H3K27me3. Nat Commun 11, 3140, doi:10.1038/s41467-020-16907-0 (2020).

33. Blondel, V. D., Guillaume, J. L., Lambiotte, R. & Lefebvre, E. Fast unfolding of communities in large networks. J Stat Mech-Theory E, doi:Artn P10008 10.1088/1742-5468/2008/10/P10008 (2008).

34. Hao, Y. et al. Integrated analysis of multimodal single-cell data. Cell 184, 3573–3587 e3529, doi:10.1016/j.cell.2021.04.048 (2021).

35. Camp, J. G. et al. Human cerebral organoids recapitulate gene expression programs of fetal neocortex development. Proc Natl Acad Sci U S A 112, 15672–15677, doi:10.1073/pnas.1520760112 (2015).

36. Fleck, J. S. et al. Inferring and perturbing cell fate regulomes in human brain organoids. Nature, doi:10.1038/s41586-022-05279-8 (2022).

37. Nowakowski, T. J. et al. Spatiotemporal gene expression trajectories reveal developmental hierarchies of the human cortex. Science 358, 1318–1323, doi:10.1126/science.aap8809 (2017).

38. Kanton, S. et al. Organoid single-cell genomic atlas uncovers human-specific features of brain development. Nature 574, 418–422, doi:10.1038/s41586-019-1654-9 (2019).

39. Dougherty, J. D., Schmidt, E. F., Nakajima, M. & Heintz, N. Analytical approaches to RNA profiling data for the identification of genes enriched in specific cells. Nucleic Acids Res 38, 4218–4230, doi:10.1093/nar/gkq130 (2010).

40. Batiuk, M. Y. et al. Identification of region-specific astrocyte subtypes at single cell resolution. Nat Commun 11, 1220, doi:10.1038/s41467-019-14198-8 (2020).

41. Nomura, Y. & Segawa, M. Anatomy of Rett syndrome. Am J Med Genet Suppl 1, 289–303, doi:10.1002/ajmg.1320250529 (1986).

42. Yildirim, M. et al. Label-free three-photon imaging of intact human cerebral organoids for tracking early events in brain development and deficits in Rett syndrome. Elife 11, doi:10.7554/eLife.78079 (2022).

43. Samarasinghe, R. A. et al. Identification of neural oscillations and epileptiform changes in human brain organoids. Nat Neurosci 24, 1488–1500, doi:10.1038/s41593-021-00906-5 (2021).

44. Sun, Y. et al. Loss of MeCP2 in immature neurons leads to impaired network integration. Hum Mol Genet 28, 245–257, doi:10.1093/hmg/ddy338 (2019).

45. Smrt, R. D. et al. Mecp2 deficiency leads to delayed maturation and altered gene expression in hippocampal neurons. Neurobiol Dis 27, 77–89, doi:10.1016/j.nbd.2007.04.005 (2007).

46. Mellios, N. et al. MeCP2-regulated miRNAs control early human neurogenesis through differential effects on ERK and AKT signaling. Mol Psychiatry 23, 1051–1065, doi:10.1038/mp.2017.86 (2018).

47. Morelli, K. H. et al. MECP2-related pathways are dysregulated in a cortical organoid model of myotonic dystrophy. Sci Transl Med 14, eabn2375, doi:10.1126/scitranslmed.abn2375 (2022).

48. Gomes, A. R. et al. Modeling Rett Syndrome With Human Patient-Specific Forebrain Organoids. Front Cell Dev Biol 8, 610427, doi:10.3389/fcell.2020.610427 (2020).

49. Valles, S. L. et al. Function of Glia in Aging and the Brain Diseases. Int J Med Sci 16, 1473–1479, doi:10.7150/ijms.37769 (2019).

50. Bernaus, A., Blanco, S. & Sevilla, A. Glia Crosstalk in Neuroinflammatory Diseases. Front Cell Neurosci 14, 209, doi:10.3389/fncel.2020.00209 (2020).

51. Mayegowda, S. B. & Thomas, C. Glial pathology in neuropsychiatric disorders: a brief review. J Basic Clin Physiol Pharmacol 30, doi:10.1515/jbcpp-2018-0120 (2019).

52. Maezawa, I., Swanberg, S., Harvey, D., LaSalle, J. M. & Jin, L. W. Rett syndrome astrocytes are abnormal and spread MeCP2 deficiency through gap junctions. J Neurosci 29, 5051–5061, doi:10.1523/JNEUROSCI.0324-09.2009 (2009).

53. Ehinger, Y. et al. Analysis of Astroglial Secretomic Profile in the Mecp2-Deficient Male Mouse Model of Rett Syndrome. Int J Mol Sci 22, doi:10.3390/ijms22094316 (2021).

54. Norris, R. P. Transfer of mitochondria and endosomes between cells by gap junction internalization. Traffic 22, 174–179, doi:10.1111/tra.12786 (2021).

55. Liu, D. et al. Intercellular mitochondrial transfer as a means of tissue revitalization. Signal Transduct Target Ther 6, 65, doi:10.1038/s41392-020-00440-z (2021).

56. Saywell, V. et al. Brain magnetic resonance study of Mecp2 deletion effects on anatomy and metabolism. Biochem Biophys Res Commun 340, 776–783, doi:10.1016/j.bbrc.2005.12.080 (2006).

57. De Filippis, B. et al. Mitochondrial free radical overproduction due to respiratory chain impairment in the brain of a mouse model of Rett syndrome: protective effect of CNF1. Free Radic Biol Med 83, 167–177, doi:10.1016/j.freeradbiomed.2015.02.014 (2015).

58. Valenti, D. et al. Stimulation of the brain serotonin receptor 7 rescues mitochondrial dysfunction in female mice from two models of Rett syndrome. Neuropharmacology 121, 79–88, doi:10.1016/j.neuropharm.2017.04.024 (2017).

59. Kriaucionis, S. et al. Gene expression analysis exposes mitochondrial abnormalities in a mouse model of Rett syndrome. Mol Cell Biol 26, 5033–5042, doi:10.1128/MCB.01665-05 (2006).

60. Grosser, E. et al. Oxidative burden and mitochondrial dysfunction in a mouse model of Rett syndrome. Neurobiol Dis 48, 102–114, doi:10.1016/j.nbd.2012.06.007 (2012).

61. Muller, M. & Can, K. Aberrant redox homoeostasis and mitochondrial dysfunction in Rett syndrome. Biochem Soc Trans 42, 959–964, doi:10.1042/BST20140071 (2014).

62. Toloe, J., Mollajew, R., Kugler, S. & Mironov, S. L. Metabolic differences in hippocampal ’Rett’ neurons revealed by ATP imaging. Mol Cell Neurosci 59, 47–56, doi:10.1016/j.mcn.2013.12.008 (2014).

63. Bebensee, D. F., Can, K. & Muller, M. Increased Mitochondrial Mass and Cytosolic Redox Imbalance in Hippocampal Astrocytes of a Mouse Model of Rett Syndrome: Subcellular Changes Revealed by Ratiometric Imaging of JC-1 and roGFP1 Fluorescence. Oxid Med Cell Longev 2017, 3064016, doi:10.1155/2017/3064016 (2017).

64. Li, J. et al. Conservation and divergence of vulnerability and responses to stressors between human and mouse astrocytes. Nat Commun 12, 3958, doi:10.1038/s41467-021-24232-3 (2021).

65. Li, C. H. et al. MeCP2 links heterochromatin condensates and neurodevelopmental disease. Nature 586, 440–444, doi:10.1038/s41586-020-2574-4 (2020).

66. Muffat, J. et al. Human induced pluripotent stem cell-derived glial cells and neural progenitors display divergent responses to Zika and dengue infections. Proc Natl Acad Sci U S A 115, 7117–7122, doi:10.1073/pnas.1719266115 (2018).

67. Fernandopulle, M. S. et al. Transcription Factor-Mediated Differentiation of Human iPSCs into Neurons. Curr Protoc Cell Biol 79, e51, doi:10.1002/cpcb.51 (2018).

68. Rappsilber, J., Mann, M. & Ishihama, Y. Protocol for micro-purification, enrichment, pre-fractionation and storage of peptides for proteomics using StageTips. Nat Protoc 2, 1896–1906, doi:10.1038/nprot.2007.261 (2007).

69. Rath, S. et al. MitoCarta3.0: an updated mitochondrial proteome now with sub-organelle localization and pathway annotations. Nucleic Acids Res 49, D1541–D1547, doi:10.1093/nar/gkaa1011 (2021).

70. Schulte, F., Hasturk, H. & Hardt, M. Mapping Relative Differences in Human Salivary Gland Secretions by Dried Saliva Spot Sampling and nanoLC-MS/MS. Proteomics 19, e1900023, doi:10.1002/pmic.201900023 (2019).

71. Tapia, J. C. et al. High-contrast en bloc staining of neuronal tissue for field emission scanning electron microscopy. Nat Protoc 7, 193–206, doi:10.1038/nprot.2011.439 (2012).

72. Lam, J. et al. A Universal Approach to Analyzing Transmission Electron Microscopy with ImageJ. Cells 10, doi:10.3390/cells10092177 (2021).

73. He, Z., Brazovskaja, A., Ebert, S., Camp, J. G. & Treutlein, B. CSS: cluster similarity spectrum integration of single-cell genomics data. Genome Biol 21, 224, doi:10.1186/s13059-020-02147-4 (2020).

74. Stuart, T., Srivastava, A., Madad, S., Lareau, C. A. & Satija, R. Single-cell chromatin state analysis with Signac. Nat Methods 18, 1333–1341, doi:10.1038/s41592-021-01282-5 (2021).

75. Love, M. I., Huber, W. & Anders, S. Moderated estimation of fold change and dispersion for RNA-seq data with DESeq2. Genome Biol 15, 550, doi:10.1186/s13059-014-0550-8 (2014).

76. Stark, R. & Brown, G.

77. Subramanian, A. et al. Gene set enrichment analysis: a knowledge-based approach for interpreting genome-wide expression profiles. Proc Natl Acad Sci U S A 102, 15545–15550, doi:10.1073/pnas.0506580102 (2005).

78. de Hoon, M. J., Imoto, S., Nolan, J. & Miyano, S. Open source clustering software. Bioinformatics 20, 1453–1454, doi:10.1093/bioinformatics/bth078 (2004).

